# Multi-omic integration reveals dynamic changes in human placental metabolism across gestation

**DOI:** 10.1101/2025.10.22.683996

**Authors:** Mariana Parenti, Sam Rosen, Maya Pettes, Danijel Djukovic, Daniel Raftery, Ian A. Glass, Birth Defects Research Laboratory (BDRL), Kathryn J. Gray, Alison G. Paquette, Stephen A. McCartney

## Abstract

**Objectives:** Metabolic demands of the developing conceptus are highly dynamic during pregnancy. While placental metabolism has been well described at term and in cell lines, changes in the placental metabolome during development remains understudied. We investigated the placental metabolome, metabolite trajectories, and altered pathways across trimesters in normal human pregnancy by integrating metabolomic and transcriptomic data.

**Methods:** Targeted aqueous metabolomic profiling of 372 metabolites was conducted on placental biopsies from samples collected in the first (*n*=12), second (*n*=13), and third (*n*=11) trimesters of normal pregnancy using liquid chromatography–tandem mass spectrometry.

Robust linear models identified differentially abundant metabolites across trimesters in models adjusted for fetal sex and total protein. We conducted pathway analysis using a human metabolic reconstruction. To further aid in biological interpretation, we leveraged publicly available transcriptomics data to conduct pathway-level multi-omic integration throughout gestation.

**Results:** Samples clustered by trimester in principal component analysis and we identified 5 metabolite trajectories. Out of 193 detectable metabolites, 149 (77%) differed by trimester (FDR<0.05). Using pathway-level multi-omic integration, pathways involved in extracellular transport, and pyruvate, amino acid, NAD, and membrane lipid metabolism are up-regulated in the second trimester compared to the first. In the late third trimester, pathways involved in amino acid metabolism, redox balance, mitochondrial transport, and biomolecule synthesis were down-regulated compared to second trimester.

**Conclusions:** Placental metabolite abundances change substantially across gestation and integration with metabolic gene expression provides insight into dynamic metabolic function during pregnancy. Observed pathway-level changes potentially reflect the metabolic response to invading maternal circulation in the first-to-second trimester transition, as well as changing maternal and fetal metabolic requirements. Differences observed at term may reflect placental senescence and preparation for parturition. These data can inform other molecular analyses of the placenta by providing enhanced resolution of metabolic changes across pregnancy.

## 1. INTRODUCTION

The placenta is the first organ to develop and plays critical roles in nutrient transport and resource allocation to the developing fetus as well as endocrine regulation of maternal and fetal metabolism and immune adaptation (*1–4*). To support the fetus, placental extravillous trophoblasts invade and remodel the uterine spiral arteries during in early pregnancy, allowing the placenta to access maternal circulation at the end of the first trimester (*1*). This transition from histotrophic to hemotrophic nutrition corresponds to changes in oxygen tension and nutrient availability, which supports a shift towards oxidative metabolism in later gestation. This influx of nutrients and oxygen supports placental and fetal growth through the rest of gestation. The placenta continues to grow, increasing in surface area for nutrient exchange and vascularization of the villus tree (*5*). Moreover, the placenta secretes hormones that influence maternal metabolism to ensure increased glucose and lipids are available for the developing fetus (*4*). Likewise, implantation and placentation, fetal growth, and parturition represent distinct developmental phases characterized by unique immune environments (*6*). The first trimester is associated with inflammation during implantation and placental invasion, while the phase of fetal growth starting in the second trimester is associated with an anti-inflammatory environment. However, the induction of pro-inflammatory signaling is critical for labor initiation and parturition (*6*). Thus, placental biology and function must be dynamic and responsive to changing cues and needs.

The placenta is a highly metabolic organ, and consumes 40% of the oxygen and 30% of the glucose it takes up from maternal circulation (*7, 8*). Metabolites provide needed substrates for energy metabolism; DNA, RNA, and protein synthesis; and signals and epigenetic modifications that influence gene expression (*9–11*). Little is known about the human placental metabolome across normal gestation, and most studies focus on term samples or pathological phenotypes like preeclampsia and fetal growth restriction (*12*). Placental dysfunction in preeclampsia and fetal growth restriction is associated with metabolic disruption in lipid and phospholipid metabolism, urea and nitrogen metabolism, and glutamate and branched-chain amino acid metabolism (*13–16*). We conducted a temporal analysis of changes in metabolic genes in placental and other maternal organs in mice, which revealed substantial differences in genes involved in glycophospholipid and glycosoaminoglycan biosynthesis, as well as inositol phosphate metabolism in the placenta between gestational day (GD)10 compared to GD15 and GD19 (*17*), corresponding to transition from histotrophic to hemotrophic nutrition and the mature placenta preparing for parturition (*18, 19*). This analysis pinpoints dynamic changes in placental metabolism, but it is challenging to extrapolate these metabolic changes to humans.

Systems biology provides a framework for understanding how different biological components (such as transcripts, proteins, and metabolites) interact within a system. High-throughput ‘omics techniques to comprehensively quantify transcripts, metabolites, or other biomolecular species provide a snapshot of the sample. Moreover, integrating multiple types of ‘omic data provide complementary information that can be used to resolve underlying biology (*20, 21*). Several studies of the placental transcriptome across gestation have demonstrated substantial changes in placental biology, including increasing expression of genes involved in transport, oxidative metabolism, steroid and peptide hormone synthesis, and vasculogenesis (*22–25*). Here, we investigate the placental metabolome and metabolite trajectories across normal human pregnancy and leverage publicly available transcriptomics data to conduct multi-omics integration at the pathway level to characterize placental metabolism across normal pregnancy.

## 2. METHODS

This study was approved by University of Washington Human Studies Division (UW HSD), IRB# STUDY00019253. First and second trimester placental tissues were collected in RPMI by the Birth Defects Research Laboratory (BDRL) from elective terminations from normal pregnancies without aneuploidy under UW HSD IRB# STUDY00000380. Placenta tissues at the end of gestation (third trimester) were collected by the Washington Pregnancy Biorepository (WPR) under UW HSD IRB# STUDY00010803 at term (37-0/7 – 40-6/7 weeks’ gestation) from participants recruited from the University of Washington Medical Center. In this study we included WPR participants who delivered by scheduled C-section without labor. Exclusion criteria included pregnancy complications such as multiple gestation, pre- or gestational diabetes, preeclampsia, preterm labor, or placental abnormalities as well as fetal aneuploidy and maternal medical complications including diabetes, cancer, and autoimmune disease treated with immune suppressants.

### 2.1. Sample Collection and Preparation

We used sterile instruments to collect 0.5cm-depth biopsies from multiple sites at the uterine-facing side of the placenta, which included both decidua basalis as well as placental villi in order to include both maternal and fetal tissues. Tissue biopsies were flash frozen in the vapor phase of liquid nitrogen and stored at -80°C. For sample preparation for metabolomics analysis, biopsies were thawed and 50mg of placental tissue was transferred to a microcentrifuge tube and homogenized.

Aqueous metabolites for targeted LC-MS profiling were extracted using a protein precipitation method as described elsewhere (*26, 27*). Placenta tissue samples were first homogenized in 200 µL purified deionized water at 4 °C, and then 800 µL of cold methanol containing the reference internal standards 124 µM 6C13-glucose and 25.9 µM 2C13-glutamate was added. The internal standards were used to monitor sample preparation. Afterwards, samples were vortexed, stored for 30 min at -20 °C, sonicated in an ice bath for 10 min, centrifuged for 15 min at 14,000 rpm and 4 °C, and then 600 µL of supernatant was collected from each sample. The total amount of protein (µg) in each sample was determined by the bicinchoninic acid (BCA) assay to enable adjustment for sample mass. Lastly, recovered supernatants were dried on a SpeedVac and reconstituted in 0.5 mL of LC-matching solvent containing 17.8 µM 2C13-tyrosine and 39.2 3C13-lactate as reference internal standards to monitor LC-MS performance. Samples were transferred into LC vials and placed into a temperature controlled autosampler set at 4 °C for LC-MS analysis.

### 2.2. Targeted Aqueous Metabolomics Assay

Targeted LC-MS metabolite analysis was performed on a duplex-LC-MS system composed of two Shimadzu UPLC pumps, CTC Analytics PAL HTC-xt temperature-controlled auto-sampler and AB Sciex 6500+ Triple Quadrupole MS equipped with an electrospray ionization (ESI) source (*26, 27*). UPLC pumps were connected to the autosampler in parallel and could perform two chromatography separations in parallel independently from each other. Each sample was injected twice on two identical analytical columns (Waters XBridge BEH Amide Premier 2.1 x 150mm, Waters part # 186009930) performing separations in hydrophilic interaction liquid chromatography (HILIC) mode. While one column performed separation for MS data acquisition in ESI positive ionization mode, the other column was equilibrated for sample injection, chromatography separation and MS data acquisition in ESI negative mode. Each chromatography separation was 16 min (for a total analysis time of 32 min per sample), and MS data acquisition was performed in multiple-reaction-monitoring (MRM) mode. The LC-MS system was controlled using AB Sciex Analyst 1.7.1 software. The LC-MS assay targeted 372 metabolic species and 4 stable isotope labeled internal standards (376 MRMs). In addition to the study samples, two sets of quality control (QC) samples were used to monitor the assay performance and data reproducibility. One in-house QC [QC(I)] was a pooled human serum sample used to monitor system performance, and the other QC [QC(S)] consisted of pooled study samples, which was used to monitor data reproducibility. Each QC sample was injected once for every 10 study samples. All the samples were analyzed in a single batch over approximately 24 hours of non-stop data acquisition. 233 metabolites (plus four spiked standards) were measured across the study set and the median coefficient of variation (**CV**) was under 5% (based on 3 QC and measured peak areas without any signal normalization). Metabolites were relatively quantified as area-under-the-curves of the MRM transitions using AB Sciex MultiQuant 3.0.3 software.

### 2.4. Data Analysis

We retained 193 metabolites that were detected in all samples and had CV < 20% for further analysis. Metabolite abundances and total protein were standardized with mean-centering and unit-scaling. Because the placenta secretes a variety of proteins across gestation at different levels, total protein concentrations could vary with gestational age (*28, 29*). Thus, all metabolomics models were adjusted for total protein and fetal sex as covariates. We employed principal components analysis (PCA, *stats::prcomp*) to reduce dimensionality and visualize the placental metabolome in low-dimensional space. We evaluated associations between the metabolome in low dimensional space and trimester using multivariate analysis of variance (**MANOVA**, *car::Manova*) adjusted for fetal sex and total protein.

We used robust linear models with M-estimation (*MASS::rlm*) to identify differentially abundant metabolites (**DAMs**) across trimesters in models adjusted for fetal sex and total protein. To asses significance, we used robust *F* tests (*sfsmisc::f*.*robftest*) to conduct hypothesis tests and corrected for false discovery rate (**FDR**) using the Benjamini-Hochberg procedure (*30, 31*). We considered FDR<0.05 to be significant.

#### 2.4.1. Exploratory Metabolite Trajectory Clustering

We conducted a hierarchical clustering analysis of average metabolite abundances across gestation. We estimated marginal means (*emmeans::emmeans*) for each trimester in robust linear models adjusted for fetal sex and total protein. Metabolites were clustered based on Euclidean distance using Ward’s clustering criterion. We calculated silhouette width (*cluster::silhouette*) to identify the optimal number of clusters by maximizing the average silhouette width.

#### 2.4.2. Transcriptomic Analysis

We leveraged independent transcriptomic data collected in placental samples across gestation (GEO accession GSE222032) (*22*), including 12 first trimester (8-12 weeks’ gestation), 12 second trimester (15-22 week’s gestation), and 12 third trimester samples (39 weeks’ gestation) to better understand metabolic changes in the placenta. RNA-sequencing data was normalized to log counts per million (**logCPM**), filtered to remove transcripts with low mean expression (logCPM<0), constrained to metabolic genes collated within the Recon3D human metabolic reconstruction (*32*), and normalized to the trimmed means of M. Trimester-specific differences in placental gene expression for 1873 metabolic genes was evaluated in models adjusted for fetal sex using the limma-voom pipeline (*33*) and differentially expressed genes (**DEGs**) with FDR<0.05 were considered significantly differentially expressed. We further identified genes encoding rate-limiting enzymes to identify critical genes regulating metabolic pathways (*34*).

#### 2.4.3. Metabolomics-Based Pathway Analysis

Pathway analysis methods were originally developed for transcripts, and metabolomics-based pathway analysis presents unique challenges (*35*). First, metabolic pathways are compartmentalized in different tissues and metabolites might not be synthesized in the tissue under study (*35*). We address this by limiting pathways to the metabolic reactions catalyzed by placentally expressed enzymes. Second, intermediate metabolites are synthesized and consumed within a pathway, and might not be elevated within an activated pathway (*35*). We address this by identifying important metabolic products of interest and DEGs encoding rate-limiting enzymes (*34*). Third, multiple pathways converge to produce and synthesize important intermediates, which participate in multiple metabolic pathways, so the same set of DAMs might drive differences in pathways. We address this challenge below through multi-omics integration.

To conduct metabolomics-based pathway analysis, we used a genome-scale human metabolic reconstruction Recon3D (*32*) and constrained the model to metabolic reactions associated with the placentally expressed genes included in our transcriptomic analysis (GSE222032). This ensures that the analysis is limited to metabolic reactions relevant to the placenta. Within Recon3D, each reaction and its associated metabolites (identified as KEGG compounds) are categorized into pathways called molecular subsystems.

We conducted pathway analysis using the Goeman’s global test (**GGT**, *globaltest::gt*) to determine if the metabolites in a metabolic subsystem were generally regulated in the same direction (*36*). This self-contained test uses sample-level metabolite abundance data to identify pathways that are consistently upregulated or downregulated in each trimester comparison.

These models were also adjusted for total protein and fetal sex. For the trajectory analysis, we were limited to metabolite lists and conducted over-representation analysis (**ORA**) using one-sided Fisher’s exact test to determine if the metabolites assigned to a trajectory cluster were enriched in molecular subsystems (*37*). We limited the background metabolites to Recon3D metabolites measured in this analysis, as is best practice in bioinformatics (*38*). This is because nonspecific background lists bias over-representation tests away from the null (*39*). We limited the analysis to compounds that resolved to a single KEGG identifier and restricted the tests to subsystems with at least 5 metabolites in the background. To guide biological interpretation of DAMs, we considered subsystems with FDR<0.05 to be significant.

#### 2.4.4. Transcriptomics-Based Pathway Analysis

To conduct transcriptomics-based pathway analysis, we similarly constrained Recon3D network to the placentally expressed genes in our analysis as background. Mapping genes to metabolic subsystems, we identified up and down-regulated subsystems using rotational gene set testing (**ROAST**) with 9,999 rotations implemented using *limma::mroast* (*40*).

#### 2.4.5. Multi-omic Pathway Integration

Transcriptomic and metabolomic data provide complementary information about the underlying biological system (*20*). We integrated the metabolic and transcriptomic data at the pathway level using Stouffer’s *Z*-score method for pooled *p*-value meta-analysis (*20, 41*). A benefit of this approach is that it can be used to combine results from multiple tests that might not have directly comparable effect sizes, such as when conducting multi-omic integration. We considered subsystems with FDR<0.05 to be significant.

## 3. RESULTS

This metabolomics analysis included *N*=36 placental tissue samples collected throughout gestation (**Table 1**). Samples clustered by trimester based on PCA (**Figure 1A**). The first and second principal components (describing 27.0% and 15.9% of the variance, respectively) were significantly associated with trimester (*p* = 2.45 x 10^-13^, MANOVA) after adjustment for fetal sex and total protein.

**Table 1.** Clinical characteristics by trimester of collection.

|  | **First trimester (N = 12)** | **Second Trimester (N = 13)** | **Third Trimester (N = 11)** |
| --- | --- | --- | --- |
| Maternal age in years, mean (range) | 27 (20-37) | 27 (19-40) | 34 (30-38) |
| Gravidity, median (range) | 2 (1-5) | 2 (1-5) | 2 (1-9) |
| Parity, median (range) | 1 (0-4) | 1 (0-4) | 2 (0-8) |
| Non-white or Hispanic, N (%) | 8 (67%) | 7 (53%) | 5 (45%) |
| Male fetal sex, N (%) | 5 (42%) | 7 (54%) | 2 (18%) |
| Gestational age in weeks, mean (range) | 11.5 (8-13) | 20 (16-23) | 39 (37-41) |

**Figure 1.**
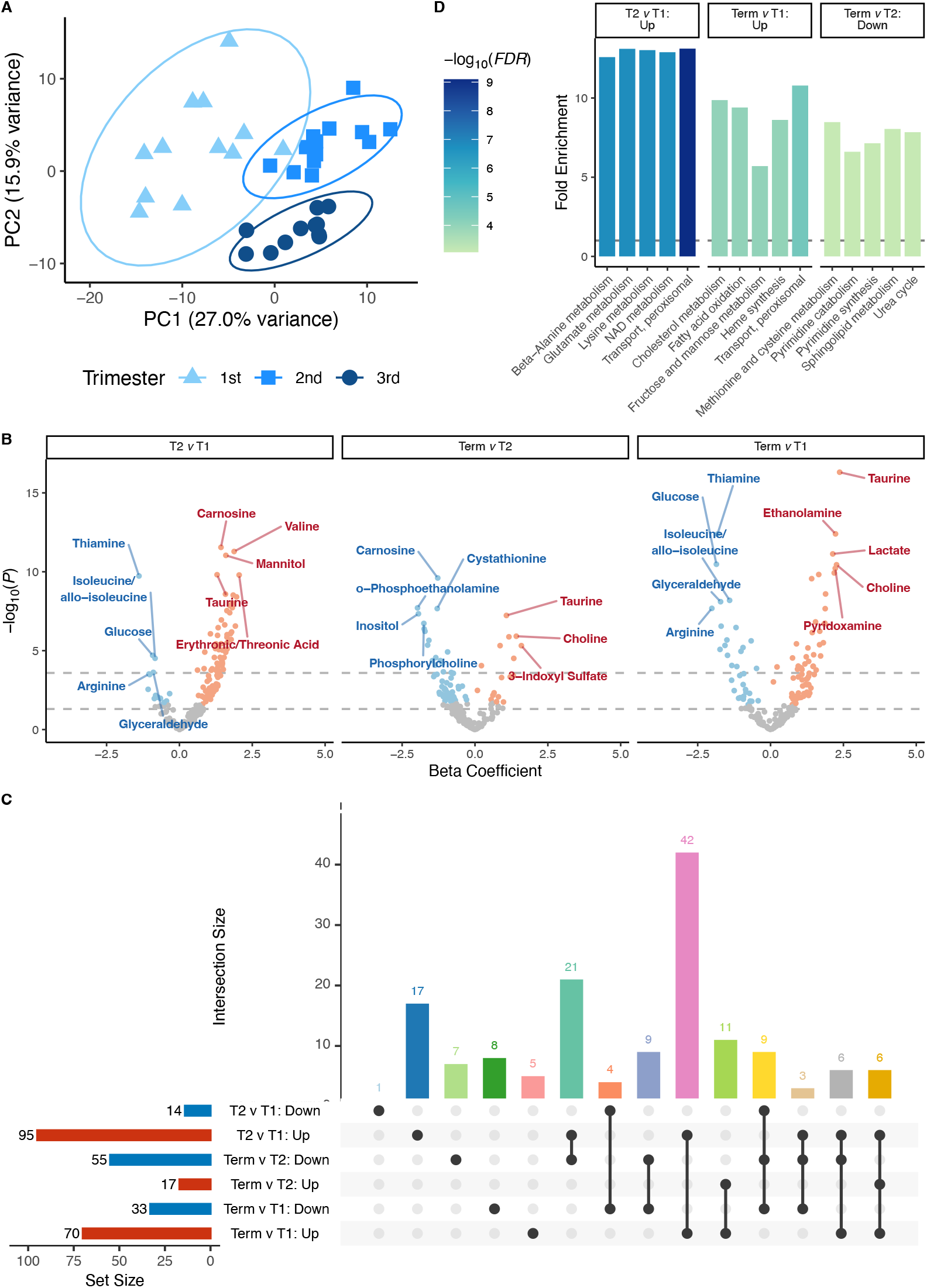
The trimester of gestation is associated with substantial changes in the placental metabolome. (**A**) Principal components analysis (PCA) reveals that samples cluster by trimester. Ellipses depict 95% confidence ellipses for each trimester. (**B**) Volcano plots showing the comparison of second trimester to first trimester (T2 v T1), third trimester at term to first trimester (Term v T1), and third trimester at term to second trimester (Term v T2). The top 5 up- and down-regulated metabolites in each comparison are labeled. (**C**) Upset plot of shared and distinct changes in placental metabolite abundance across trimesters. (D) The top 5 enriched subsystems for each comparison from directional quantitative enrichment analysis (FDR<0.05).

### 3.1. Differentially Abundant Metabolites and Pathway Enrichment

Using robust linear models, we identified metabolites that were differentially abundant in each trimester (**Figure 1B**) and present the full results in **Supplementary Table 1**. In total, 149 of 193 (77%) metabolites were differentially abundant between at least 2 trimesters (**Figure 1C**). In the second trimester compared to the first trimester (**T2vT1**), we identified 95 DAMs that were more abundant in the second trimester and 14 DAMs that were less abundant in the second trimester. At term compared to the first trimester (**TermvT1**), we also identified 70 DAMs that were more abundant in the third trimester and 33 DAMs that were less abundant in the third trimester compared to the first trimester. Finally, at term compared to the second trimester (**TermvT2**), we identified 17 DAMs more abundant in the third trimester and 55 DAMs less abundant in the third trimester. We used GGT on 49 Recon3D molecular subsystems to identify molecular subsystems that were differentially regulated with FDR<0.05 between trimesters (**Figure 1D** and **Supplementary Table 2**). Compared to the first trimester, 48 subsystems were upregulated in the second trimester, and 34 subsystems were upregulated in the third trimester at term. No subsystems were downregulated in the second or third trimesters compared to the first. Compared to the second trimester, 32 subsystems were downregulated in the third trimester at term and no subsystems were significantly upregulated.

**Table 2.** Multi-omic Pathway-Level Integration. Using Stouffer’s test to integrate metabolomics and transcriptomics data at the pathway level, we present significant (FDR<0.05) subsystems for each contrast. To support interpretation, we highlight differentially expressed genes (DEGs) encoding rate-limiting enzymes (RLE) and differentially abundant metabolites (DAMs) that are directionally regulated with the pathway. FE, fold enrichment; Gene prop., gene proportion regulated in the subsystem direction; GGT, Goeman’s global test; ROAST, rotational gene set testing.

| Contrast | Direction | Molecular Subsystem | Metabolomics |  |  | Transcriptomics |  |  | Integration FDR |
| --- | --- | --- | --- | --- | --- | --- | --- | --- | --- |
|  |  |  | FE | GGT *P* | DAMs in subsystem | Gene prop. | ROAST *P* | RLE DEGs |  |
| T2vT1 | Up | Aminosugar metabolism | 7.6 | 6.27E-05 | ADP, Glutamic acid, NAD, UDP-GlcNAc | 0.39 | 0.0003 |  | 1.72E-06 |
| T2vT1 | Up | Arachidonic acid metabolism | 10.2 | 2.70E-06 | AMP, NAD, Oxidized Glutathione | 0.22 | 0.2947 | *ALOX5, PTGS2* | 3.94E-04 |
| T2vT1 | Up | Beta-Alanine metabolism | 12.6 | 1.60E-08 | AMP, betaAlanine, Carnosine, gamma-Aminobutyrate, Glutamic acid, NAD | 0.21 | 0.9929 |  | 4.31E-04 |
| T2vT1 | Up | Bile acid synthesis | 8.5 | 1.33E-04 | ADP, AMP, GMP, NAD | 0.29 | 0.3911 | *HSD3B2, STS* | 2.82E-03 |
| T2vT1 | Up | Glycerophospholipid metabolism | 11.3 | 1.79E-07 | Acetylcholine, ADP, Choline, Ethanolamine, Inositol, o-Phosphoethanolamine, Phosphorylcholine, S-Adenosylmethionine (SAM), SAH | 0.31 | 0.0023 |  | 2.40E-07 |
| T2vT1 | Up | Histidine metabolism | 11.2 | 4.03E-07 | 1-Methylimidazole Acetate, Glutamic acid, Glycine, NAD, S-Adenosylmethionine (SAM), SAH | 0.25 | 0.8462 |  | 7.34E-04 |
| T2vT1 | Up | N-glycan synthesis | 1.9 | 1.16E-01 | UDP-GlcNAc, UDP-Glucose | 0.36 | 0.0497 |  | 2.99E-02 |
| T2vT1 | Up | NAD metabolism | 12.9 | 9.94E-09 | 1-Methylnicotinamide, ADP, AMP, Glutamic acid, NAD, Niacinamide, S-Adenosylmethionine (SAM), SAH | 0.22 | 0.2066 | *NT5E* | 7.27E-06 |
| T2vT1 | Up | Nucleotide interconversion | 6.6 | 1.00E-03 | ADP, AMP, Aspartic Acid, Fumaric Acid, Glutamic acid, GMP, Hypoxanthine, NAD, Uracil, Xanthine | 0.28 | 0.6885 | *NT5E, RRM2B* | 2.29E-02 |
| T2vT1 | Up | Protein degradation | 5.1 | 1.04E-02 | ADP, Alanine, Aspartic Acid, Glutamic acid, Glycine, Lysine, Phenylalanine, Proline, Threonine, Tyrosine, Valine | 0.56 | 0.0028 |  | 4.41E-04 |
| T2vT1 | Up | Pyrimidine catabolism | 5.8 | 1.12E-04 | ADP, Glutamic acid, Thymine, Uracil | 0.38 | 0.0954 | *NT5E* | 4.41E-04 |
| T2vT1 | Up | Pyruvate metabolism | 10.2 | 2.16E-07 | ADP, Lactate, NAD | 0.24 | 0.9210 |  | 7.34E-04 |
| T2vT1 | Up | Sphingolipid metabolism | 9.2 | 2.61E-07 | ADP, o-Phosphoethanolamine, Phosphorylcholine, UDP-GlcNAc, UDP-Glucose | 0.32 | 0.0537 |  | 6.68E-06 |
| T2vT1 | Up | Starch and sucrose metabolism | 4.1 | 3.41E-03 | NAD, UDP-Glucose | 0.23 | 0.6889 | *GYS2* | 4.25E-02 |
| T2vT1 | Up | Transport, endoplasmic reticular | 4.8 | 1.52E-03 | Acetylcarnitine, ADP, UDP-GlcNAc | 0.29 | 0.4044 | *CPT1A, PTGS2* | 1.24E-02 |
| T2vT1 | Up | Transport, extracellular | 8.1 | 1.99E-04 | Acetylcarnitine, Acetylcholine, ADP, Alanine, AMP, Aspartic Acid, betaAlanine, Carnosine, Choline, Creatine, Epinephrine, gamma-Aminobutyrate, Glutamic acid, Glycine, | 0.29 | 0.1428 |  | 8.41E-04 |
|  |  |  | FE | GGT *P* | DAMs in subsystem | Gene prop. | ROAST *P* | RLE DEGs |  |
|  |  |  |  |  | Hypoxanthine, Inositol, L-Kynurenine, Lactate, Lysine, N-Ac-Methionine, NAD, Phenylalanine, Proline, SAH, Taurine, Threonine, Thymine, Tyrosine, Uracil, Urate, Valine |  |  |  |  |
| T2vT1 | Up | Transport, golgi apparatus | 5.2 | 1.86E-03 | ADP, GMP, Sorbitol, UDP-GlcNAc, UDP-Glucose | 0.25 | 0.2375 |  | 6.78E-03 |
| T2vT1 | Up | Transport, lysosomal | 7.5 | 2.82E-04 | ADP, Alanine, Carnosine, gamma-Aminobutyrate, Glycine, Hypoxanthine, Proline, Sorbitol | 0.24 | 0.2616 |  | 2.48E-03 |
| T2vT1 | Up | Tryptophan metabolism | 10.6 | 5.79E-07 | Alanine, Glutamic acid, L-Kynurenine, NAD, S-Adenosylmethionine (SAM), SAH | 0.24 | 0.5377 | *IDO1* | 4.31E-04 |
| T2vT1 | Up | Tyrosine metabolism | 12.6 | 2.44E-08 | Epinephrine, Fumaric Acid, Glutamic acid, NAD, S-Adenosylmethionine (SAM), SAH, Tyrosine | 0.29 | 0.0618 | *STS* | 2.25E-06 |
| T2vT1 | Up | Xenobiotics metabolism | 8.4 | 1.56E-04 | ADP, AMP, NAD | 0.27 | 0.1040 |  | 5.75E-04 |
| TermvT1 | Down | Alanine and aspartate metabolism | 2.8 | 6.32E-02 | Alpha-Ketoglutaric Acid, Arginine, Asparagine, Citrulline, Glutamine | 0.55 | 0.0004 | *ASS1* | 2.79E-03 |
| TermvT1 | Down | Methionine and cysteine metabolism | 1.7 | 1.56E-01 | Alpha-Ketoglutaric Acid, Cystathionine, Cysteine, Methionine, Putrescine | 0.48 | 0.0001 | *CBS, CTH, MDH1, MDH2* | 2.79E-03 |
| TermvT1 | Down | Selenoamino acid metabolism | 1.3 | 2.50E-01 | Selenomethionine | 0.42 | 0.0005 | *CBS, CTH* | 1.27E-02 |
| TermvT1 | Down | Transport, mitochondrial | 1.8 | 1.51E-01 | Alpha-Ketoglutaric Acid, Arginine, Biotin, Citrulline, Uridine | 0.53 | 0.0001 | *ACSL1, CPT1A, SLC27A2, SLC27A5* | 2.79E-03 |
| TermvT1 | Up | Aminosugar metabolism | 6.3 | 2.39E-04 | ADP, NAD, NADH | 0.46 | 0.0014 | *GFPT1, GFPT2, GNE* | 1.43E-05 |
| TermvT1 | Up | Glycerophospholipid metabolism | 6.7 | 2.09E-04 | ADP, Choline, Ethanolamine, Glycerophosphocholine, SAH | 0.52 | 0.0001 | *GPAT3, LIPE, LPL, PCYT1B, PIP5K1A, PIP5K1B, PLA2G12A, PLA2G5, PLA2G6, PLPP2, PLPP3* | 1.24E-06 |
| TermvT1 | Up | N-glycan synthesis | 6.8 | 8.25E-05 |  | 0.61 | 0.0001 |  | 8.66E-07 |
| TermvT1 | Up | NAD metabolism | 8.5 | 3.38E-05 | ADP, NAD, NADH, Niacinamide, SAH | 0.48 | 0.0377 | *NT5C2, NT5E* | 8.61E-05 |
| TermvT1 | Up | Nucleotide interconversion | 2.4 | 1.01E-01 | Adenine, ADP, Fumaric Acid, Hypoxanthine, NAD, NADH, Uracil | 0.42 | 0.0485 | *ADK, NT5C2, NT5E, RRM2B, UCK1, UCK2, UCKL1, XDH* | 4.26E-02 |
| TermvT1 | Up | Pyrimidine catabolism | 1.3 | 2.55E-01 | ADP, Uracil | 0.52 | 0.0001 | *DPYD, NT5C2, NT5E* | 2.01E-03 |
| TermvT1 | Up | Sphingolipid metabolism | 3.4 | 1.75E-02 | ADP | 0.50 | 0.0001 | *PLPP2, PLPP3, SPTLC1, SPTLC3* | 8.61E-05 |
| TermvT1 | Up | Transport, endoplasmic reticular | 4.4 | 1.48E-03 | Acetylcarnitine, ADP | 0.42 | 0.2332 | *CPT1B, PTGS2* | 8.73E-03 |
| TermvT1 | Up | Transport, golgi apparatus | 2.4 | 6.63E-02 | ADP | 0.61 | 0.0001 |  | 3.52E-04 |
|  |  |  | FE | GGT *P* | DAMs in subsystem | Gene prop. | ROAST *P* | RLE DEGs |  |
| TermvT1 | Up | Transport, lysosomal | 2.3 | 8.88E-02 | Adenine, ADP, Alanine, gamma-Aminobutyrate, Glucuronate, Hypoxanthine, Proline | 0.55 | 0.0001 |  | 4.60E-04 |
| TermvT1 | Up | Tyrosine metabolism | 7.5 | 9.61E-05 | Fumaric Acid, NAD, NADH, SAH, Tyrosine | 0.53 | 0.0001 | *ADH6, GSTK1, STS* | 8.66E-07 |
| TermvT1 | Up | Xenobiotics metabolism | 8.6 | 3.00E-05 | ADP, Glucuronate, NAD, NADH | 0.35 | 0.4754 | *ACOX1, ADH6, CYP19A1, GSTA3, GSTK1, MGST2, XDH* | 2.64E-03 |
| TermvT2 | Down | Alanine and aspartate metabolism | 8.1 | 1.22E-04 | Arginine, Asparagine, Glutamine, N-Acetyl-Aspartate (NAA) | 0.55 | 0.0004 | *ASS1* | 1.43E-06 |
| TermvT2 | Down | Aminoacyl-tRNA biosynthesis | 8.5 | 1.38E-04 | Arginine, Asparagine, Cysteine, Glutamine, Glycine, Histidine, Leucine /D-Norleucine, Lysine, Methionine, Phenylalanine, Proline, Serine, Threonine | 0.59 | 0.0001 |  | 1.11E-06 |
| TermvT2 | Down | Aminosugar metabolism | 3.8 | 7.74E-03 | Glutamine | 0.43 | 0.5525 | *GCK, HK2, HK3* | 4.25E-02 |
| TermvT2 | Down | Arachidonic acid metabolism | 1.8 | 1.50E-01 |  | 0.48 | 0.0010 | *CYP2S1, HADH, PTGS1, SLC27A2* | 1.98E-03 |
| TermvT2 | Down | Arginine and proline metabolism | 7.3 | 1.98E-04 | Arginine, Hydroxyproline, Proline, Putrescine, S-Adenosylmethionine (SAM) | 0.62 | 0.0002 | *ALDH1A1, ALDH1A2, ALDH1B1, ALDH2, ALDH3A2, ALDH7A1, ALDH9A1, NOS2, ODC1, SAT2* | 1.37E-06 |
| TermvT2 | Down | Beta-Alanine metabolism | 6.5 | 2.50E-04 | betaAlanine, Carnosine, gamma-Aminobutyrate, Histidine | 0.58 | 0.0001 | *ALDH1A1, ALDH1A2, ALDH1B1, ALDH2, ALDH3A2, ALDH7A1, ALDH9A1* | 1.11E-06 |
| TermvT2 | Down | Butanoate metabolism | 0.3 | 8.19E-01 |  | 0.73 | 0.0005 |  | 1.82E-02 |
| TermvT2 | Down | Fructose and mannose metabolism | 2.7 | 5.83E-02 | Glucose, Mannose | 0.65 | 0.0001 | *FBP1, GCK, HK2, HK3* | 1.48E-04 |
| TermvT2 | Down | Glutamate metabolism | 6.3 | 9.28E-04 | gamma-Aminobutyrate, Glutamine, Pyruvate | 0.61 | 0.0001 | *ALDH1B1, ALDH2, ALDH3A2, ALDH7A1, ALDH9A1* | 2.12E-06 |
| TermvT2 | Down | Glutathione metabolism | 5.3 | 1.84E-03 | Cysteine, Glycine, Pyroglutamic Acid | 0.47 | 0.0001 | *GGT7* | 3.17E-06 |
| TermvT2 | Down | Glycine, serine, alanine, and threonine metabolism | 5.5 | 1.15E-03 | Aminolevulinate, Glycine, Methionine, Phosphoserine, Serine, Threonine | 0.67 | 0.0001 | *ALDH7A1* | 2.12E-06 |
| TermvT2 | Down | Glycolysis/gluconeogenesis | 2.2 | 1.01E-01 | Glucose, Pyruvate | 0.54 | 0.0001 | *ADH5, ALDH1A1, ALDH1A2, ALDH1B1, ALDH2, ALDH3A2, ALDH7A1, ALDH9A1, FBP1, GCK, HK2, HK3, PCK2, PFKM, PFKP, PKM* | 2.77E-04 |
| TermvT2 | Down | Heme synthesis | 1.8 | 1.29E-01 | Aminolevulinate | 0.54 | 0.0001 | *CPOX, HMBS* | 3.70E-04 |
|  |  |  | FE | GGT *P* | DAMs in subsystem | Gene prop. | ROAST *P* | RLE DEGs |  |
| TermvT2 | Down | Histidine metabolism | 3.5 | 1.18E-02 | Glycine, S-Adenosylmethionine (SAM) | 0.50 | 0.0118 | *ALDH1A1, ALDH1A2, ALDH1B1, ALDH2, ALDH3A2, ALDH7A1, ALDH9A1* | 9.36E-04 |
| TermvT2 | Down | Methionine and cysteine metabolism | 8.5 | 2.77E-05 | Cystathionine, Cysteine, Methionine, N-Ac-Methionine, Putrescine, Pyruvate, S-Adenosylmethionine (SAM), Serine | 0.45 | 0.0025 | *CTH* | 2.12E-06 |
| TermvT2 | Down | Pentose phosphate pathway | 3.7 | 2.14E-02 | Erythrose-4-Phosphate, Ribulose 5-Phosphate | 0.62 | 0.0001 | *G6PD, PRPS1, PRPS2* | 4.41E-05 |
| TermvT2 | Down | Protein degradation | 8.6 | 1.33E-04 | Arginine, Asparagine, Cysteine, Glutamine, Glycine, Histidine, Leucine /D-Norleucine, Lysine, Methionine, Phenylalanine, Proline, Serine, Threonine | 0.38 | 0.6560 |  | 5.85E-03 |
| TermvT2 | Down | Pyrimidine synthesis | 7.1 | 1.12E-04 | Glutamine, Pyruvate, UMP | 0.64 | 0.0002 | *CTPS1, CTPS2, DTYMK, PKM* | 1.11E-06 |
| TermvT2 | Down | Selenoamino acid metabolism | 6.3 | 1.21E-03 | Pyruvate, Serine | 0.46 | 0.0483 | *CTH* | 6.10E-04 |
| TermvT2 | Down | Starch and sucrose metabolism | 2.8 | 5.03E-02 | Glucose, UDP-Glucose, UMP | 0.50 | 0.0256 | *GYS1, PYGL, UGDH* | 6.66E-03 |
| TermvT2 | Down | Transport, mitochondrial | 4.6 | 7.16E-03 | Arginine, Lysine, Pyruvate, S-Adenosylmethionine (SAM) | 0.47 | 0.0039 | *ACSL1, CPT1A, SLC27A2* | 2.58E-04 |
| TermvT2 | Down | Tryptophan metabolism | 4.8 | 1.07E-03 | S-Adenosylmethionine (SAM) | 0.55 | 0.0001 | *ALDH1A1, ALDH1A2, ALDH1B1, ALDH2, ALDH3A2, ALDH7A1, ALDH9A1, HADH* | 2.12E-06 |
| TermvT2 | Down | Urea cycle | 7.8 | 4.57E-05 | Arginine, Asparagine, Creatine, Glycine, N-Ac-Glutamate, Phosphocreatine, Putrescine, S-Adenosylmethionine (SAM) | 0.78 | 0.0001 | *ODC1, SRM* | 8.46E-07 |
| TermvT2 | Down | Valine, leucine, and isoleucine metabolism | 4.0 | 9.39E-03 | Leucine /D-Norleucine | 0.64 | 0.0001 | *ALDH1B1, ALDH2, ALDH3A2, ALDH7A1, ALDH9A1, HADH* | 1.79E-05 |

### 3.2. Exploratory Trajectory Clustering

We conducted exploratory metabolite trajectory clustering using hierarchical clustering with Ward’s D linkage on average metabolite abundance within each trimester. We identified the optimal number of clusters using average silhouette width (*k*=5, average silhouette width=0.368). The 5 clusters depicted in **Figure 2** correspond to an increasing trajectory (cluster 1: 20 metabolites), rapidly decreasing trajectory from second to third trimester (cluster 2: 49 metabolites), rapidly increasing trajectory from the first to second trimester (cluster 3: 53 metabolites), decreasing trajectory (cluster 4: 30 metabolites), and steady trajectory (cluster 5: 41 metabolites). Cluster assignments for each metabolite are reported in **Supplementary Table 1** and visualized in **Supplementary Figures 1-5**. Using ORA with a background of 123 metabolites, we found that the 53 metabolites in cluster 3 were enriched for the *glutamate metabolism* subsystem (8/12 metabolites, FDR = 0.013). No other clusters were significantly enriched for metabolic subsystems.

**Figure 2.**
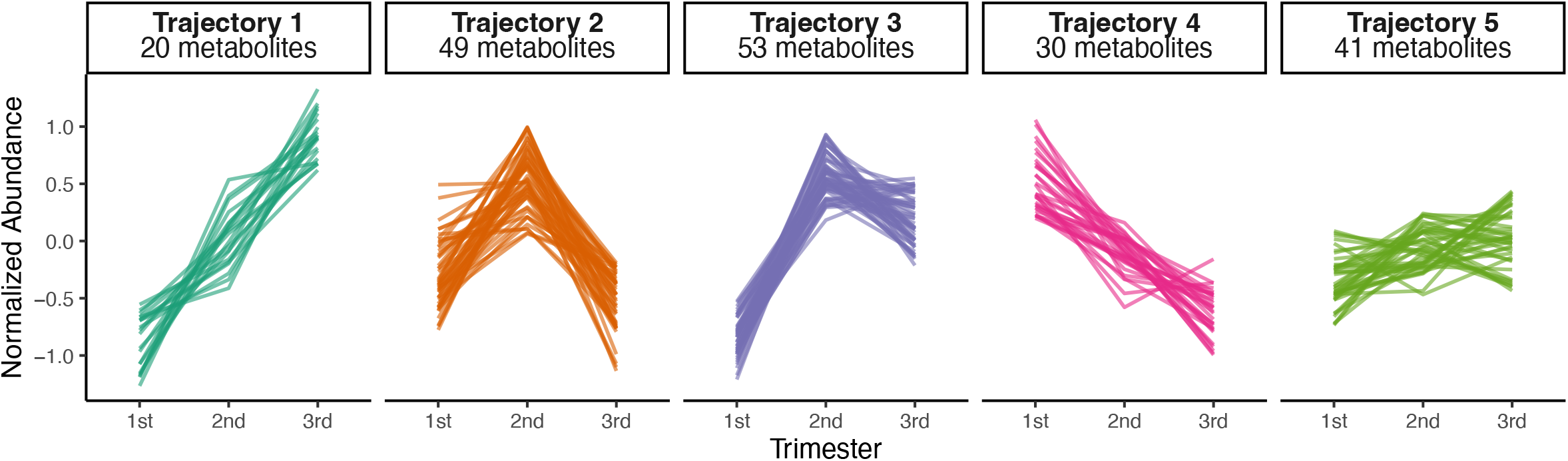
The metabolite trajectory clustering revealed 5 trajectories. Metabolites were standardized and estimated marginal means (EMM) were calculated from robust linear models adjusted for fetal sex and total protein. Each line represents a single metabolite.

### 3.3 Transcriptomics & Multi-Omics Pathway Integration

We integrated transcriptomic data (GSE222032) from metabolic genes to glean complementary information about the underlying biological changes in the placenta across gestation. We identified 1873 metabolic genes that participate in 78 subsystems and include genes encoding 108 rate-limiting enzymes (*34*). Trimester was associated with substantial differences in metabolic gene expression (**Supplementary Figure 6A-B**). Using ROAST, we identified 65 differentially regulated subsystems and report the top 5 up- and downregulated pathways for each trimester in **Supplementary Figure 6C**.

Our primary interest was understanding the metabolic changes in the placenta across gestation, so we used Stouffer’s method to integrate metabolomics and transcriptomics data (**Table 2**). Compared to the first trimester, the NAD metabolism, pyrimidine catabolism, endoplasmic reticular transport, and tyrosine metabolism were upregulated in both the second and third trimesters. Compared to the first and second trimesters, mitochondrial transport and amino acid metabolism subsystems (alanine and aspartate metabolism, methionine and cysteine metabolism, and selenoamino acid metabolism) were down-regulated. In the second trimester, tryptophan metabolism is up-regulated compared to the first and third trimesters. Compared to these second trimester, additional subsystems are down-regulated in the third trimester, including subsystems involved in protein metabolism (arginine and proline metabolism; beta-alanine metabolism; glutamate metabolism; glycine, serine, alanine, and threonine metabolism; histidine metabolism; urea cycle; valine, leucine, and isoleucine metabolism), glutathione metabolism, and pentose phosphate pathway. To better interpret these results, we have visualized results from key metabolites and rate-limiting enzymes with the integrated pathway results in a summary in **Figure 3**.

**Figure 3.**
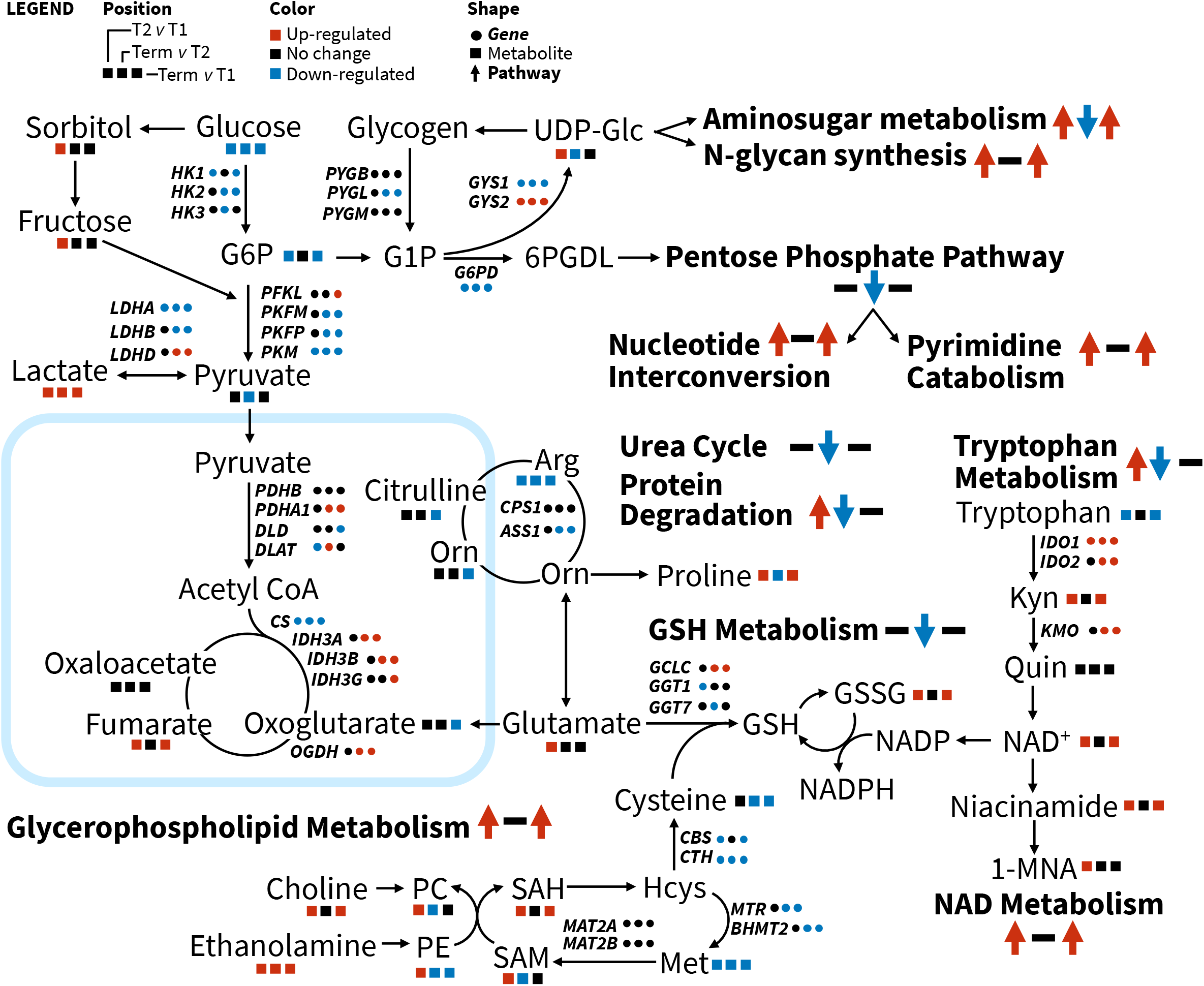
Simplified network schematic of changes in metabolite abundance and expression of key genes. Genes are denoted by HGNC gene symbols. 1-MNA, 1-Methylnicotinamide; 6PGDL, 6-Phosphonoglucono-D-lactone; Arg, arginine; G1P, glucose-1-phosphate; G6P, glucose-6-phosphate; GSH, reduced glutathione; GSSG, oxidized glutathione; Hcys, homocysteine; Kyn, kynurenine; Met, methionine; Orn, ornithine; PC, O-phosphocholine; PE, O-phosphoethanolamine; Quin, quinoline; SAH, S-adenosylhomocysteine; SAM, S-adenosylmethionine

## DISCUSSION

Our results demonstrate that the placental metabolome is dynamic across trimesters in normal pregnancy. First, we found that 149 metabolites (77%) significantly differ in relative abundance between the first, second, or third trimesters. Second, placental metabolites cluster into 5 distinct trajectories, providing insight into changing metabolic function during pregnancy. Third, we identified 49 metabolic pathways and 38 integrated pathways that were differentially regulated across gestation. This study represents an advance over prior analyses of placental metabolism by integrating independent gene expression data to gain complementary information about altered metabolic pathways.

Early in gestation before the placenta in fully functional, the conceptus experiences low oxygen tension and relies on histotrophic nutrition. Under these conditions, the placenta relies on glycolysis for energy generation (*42*). By the end of the first trimester, the placenta invades maternal circulation, resulting in a three-fold increase in oxygen tension and increased nutrient availability (*42*). This is accompanied by changes in placental gene expression and metabolism over time to support substantial changes in villus architecture, hormone synthesis, and oxidative metabolism (*42*). Here, pyruvate metabolism and extracellular transport pathways are up-regulated in the second trimester compared to the first trimester, based on our multi-omic integration. Moreover, metabolic pathways involved in protein glycosylation (amino sugar metabolism, *N*-glycan metabolism, and endoplasmic reticular transport), redox balance (NAD metabolism), membrane lipids (sphingolipid metabolism, glycerophospholipid metabolism), xenobiotic metabolism, and nucleobase metabolism (nucleotide interconversion, pyrimidine catabolism) are up-regulated in both the second and third trimesters compared to the first trimester (**Table 2**). These findings are consistent with increased placental hormone synthesis and increased oxidative metabolism after the first trimester, which are necessary for fetal growth. Upregulation of NAD metabolism could reflect the importance of reducing equivalents in oxidative metabolism together with upregulated carbohydrate metabolism in the second trimester (*43*). Protein glycosylation is a post-translational protein modification that plays an important role in cell-cell communication and immune recognition (*44*). Trophoblast glycans promote maternal-fetal immune tolerance and aberrant glycosylation has been implicated in implantation failure, early onset preeclampsia, and fetal growth restriction (*44, 45*). Metabolic pathways involved in protein glycosylation might reflect changes in maternal-fetal immunological processes in mid- and late pregnancy (*44*). Nucleobase and membrane lipid metabolism might reflect changes in metabolism to support rapid growth and proliferation of placental cells, as nucleotides and lipid bilayers make up the building blocks of new cells.

After the placenta invades maternal circulation, oxygen levels increase and peak during the second trimester (*46*). Here, the two largest clusters, cluster 2 (49 metabolites) and cluster 3 (53 metabolites), are characterized by relatively high metabolite abundances during the second trimester. Notably, cluster 3 was enriched for metabolites involved in glutamate metabolism.

Indeed, during the second trimester, 21 pathways were upregulated compared to first trimester and 24 pathways were upregulated in the second trimester compared to term in our integrated multi-omic analysis (**Table 2**). These pathways participate in protein metabolism (protein degradation, aminoacyl-tRNA biosynthesis, beta-alanine and amino acid metabolism), carbohydrate metabolism (starch and sucrose metabolism, increased mitochondrial transport, glycolysis, fructose and mannose metabolism), lipid metabolism (arachidonic acid, sphingolipid, and glycerophospholipid metabolism), and pathways related to redox balance (NAD metabolism, pentose phosphate pathway, glutathione metabolism). This pattern of high metabolism in the second trimester relative to term might reflect placental senescence at term (*47*). Finally, in the third trimester, compared to the other trimesters, pathways involved in amino acid metabolism (alanine and aspartate metabolism, selenoamino acid metabolism, and methionine and cysteine metabolism) and mitochondrial transport are downregulated. As the placenta ages, mitochondrial size and number decreases (*42*), which aligns with reduced mitochondrial transport at term compared to both first and second trimester (**Table 2**).

This work should also be interpreted in the context of its limitations. First, placentas were not collected between 23-37 weeks in our metabolomics analysis and 22-39 weeks in the transcriptomics data for practical and ethical reasons (*22*). Therefore, this analysis focused on differences between trimesters. However, it is likely that within each trimester there are also metabolic changes, particularly in the first trimester transition from histotrophic to hematologic nutrition. Thus, future studies with more precise gestational age samples are necessary to gain a full understanding of the placental metabolic transitions throughout gestation. Second, there is significant variation in frequency of decidual stromal, immune, and placental trophoblast populations throughout gestation, and metabolomics and transcriptomics analysis on bulk tissue might be confounded by differences in cell type proportion between samples and over time (*48*). Finally, pathway analysis approaches were designed for gene products, which are generally coordinately regulated and generally contribute distinct functions to pathways (*35*). Metabolites, on the other hand, are synthesized and consumed within pathways and participate in multiple biological pathways. Thus, the conservative, directional approach to metabolite pathway analysis employed here might miss real signal. However, integration with concordantly altered transcriptomic data provides an additional layer of support to these findings.

This study also has important strengths. First, this is the first temporal metabolomics study of placental tissue across gestation conducted in humans. Second, our analysis employs robust and rigorous methods, accounts for sex as a biological variable, and accounts for multiple comparisons. Third, we used a sophisticated multi-omic integration strategy to overcome limitations of metabolite-based pathway analysis.

## CONCLUSION

The placenta is a dynamic and highly metabolic organ. We report that the placental metabolome is likewise dynamic and metabolites cluster into distinct trajectories throughout gestation. Moreover, integrating metabolomics and transcriptomic data provides complementary information to better understand the underlying biological changes across gestation. Metabolomic profiling across gestation and multi-omics integration can inform other molecular analyses of the placenta by providing enhanced resolution of metabolite availability, which influences cellular function.

## Acknowledgments & Funding

We thank the UW staff and study participants for their generosity. S.A.M. is supported by the Reproductive Scientist Development Program (K12 HD000849) from both the Eunice Kennedy Shriver National Institute of Child Health & Human Development and the Burroughs Wellcome Fund as well as Doris Duke Clinical Scientist Development Award (2023-0231) from the Doris Duke Foundation and the University of Washington Diabetes, Nutrition, and Obesity Research Center Pilot and feasibility grant (P30DK017047). The Birth Defects Research Laboratory (BDRL) was supported by NIH award number 5R24HD000836 from the Eunice Kennedy Shriver National Institute of Child Health and Human Development. The content is solely the responsibility of the authors and does not necessarily represent the official views of the National Institutes of Health.

BDRL investigators include Ian A. Glass^1^, Kimberly A. Aldinger^1,2^, Dan Doherty^1^, Ian G. Phelps^1^, Jennifer C. Dempsey^1^, Kevin J. Lee^1^, and Lucinda A. Cort^1^.

^1^ University of Washington

^2^ Seattle Children’s Research Institute

