## Supplementary Materials for "Multi-omic integration reveals dynamic changes in human placental metabolism across gestation"


**Authors & Affiliations**

Mariana Parenti[1], Sam Rosen[2], Maya Pettes[2], Danijel Djukovic[3], Daniel Raftery[4], Ian A. Glass[5], Birth Defects Research Laboratory (BDRL)[5], Kathryn J. Gray[2], Alison G. Paquette[1,5,†], Stephen A. McCartney*[2,†]

[1] Center for Developmental Biology and Regenerative Medicine, Seattle Children's Research Institute, Seattle, USA
[2] Department of Obstetrics and Gynecology, University of Washington, Seattle, USA
[3] Department of Anesthesiology and Pain Medicine, University of Washington, Seattle, USA
[4] Department of Anesthesiology and Pain Medicine, University of Washington, Seattle, USA
[5] Department of Pediatrics, University of Washington, Seattle, USA
[†] These authors contributed equally


**CONTENTS**


**Supplementary Figure 1.** Cluster 1 metabolites with "increasing" trajectories.
**Supplementary Figure 2.** Cluster 2 metabolites with "decreasing into third trimester" trajectories.
**Supplementary Figure 3.** Cluster 3 metabolites with "increasing in early pregnancy" trajectories.
**Supplementary Figure 4.** Cluster 4 metabolites with "decreasing" trajectories.
**Supplementary Figure 5.** Cluster 4 metabolites with "steady" trajectories.
**Supplementary Figure 6**: Transcriptomics analysis of metabolic genes.
**Supplementary Table 1.** Differential abudance of metabolites with different trajectories in placental samples across three trimesters.
**Supplementary Table 2**. Directional Goeman's global tests (GGT) of Recon3D molecular subsystems (FDR<0.05).

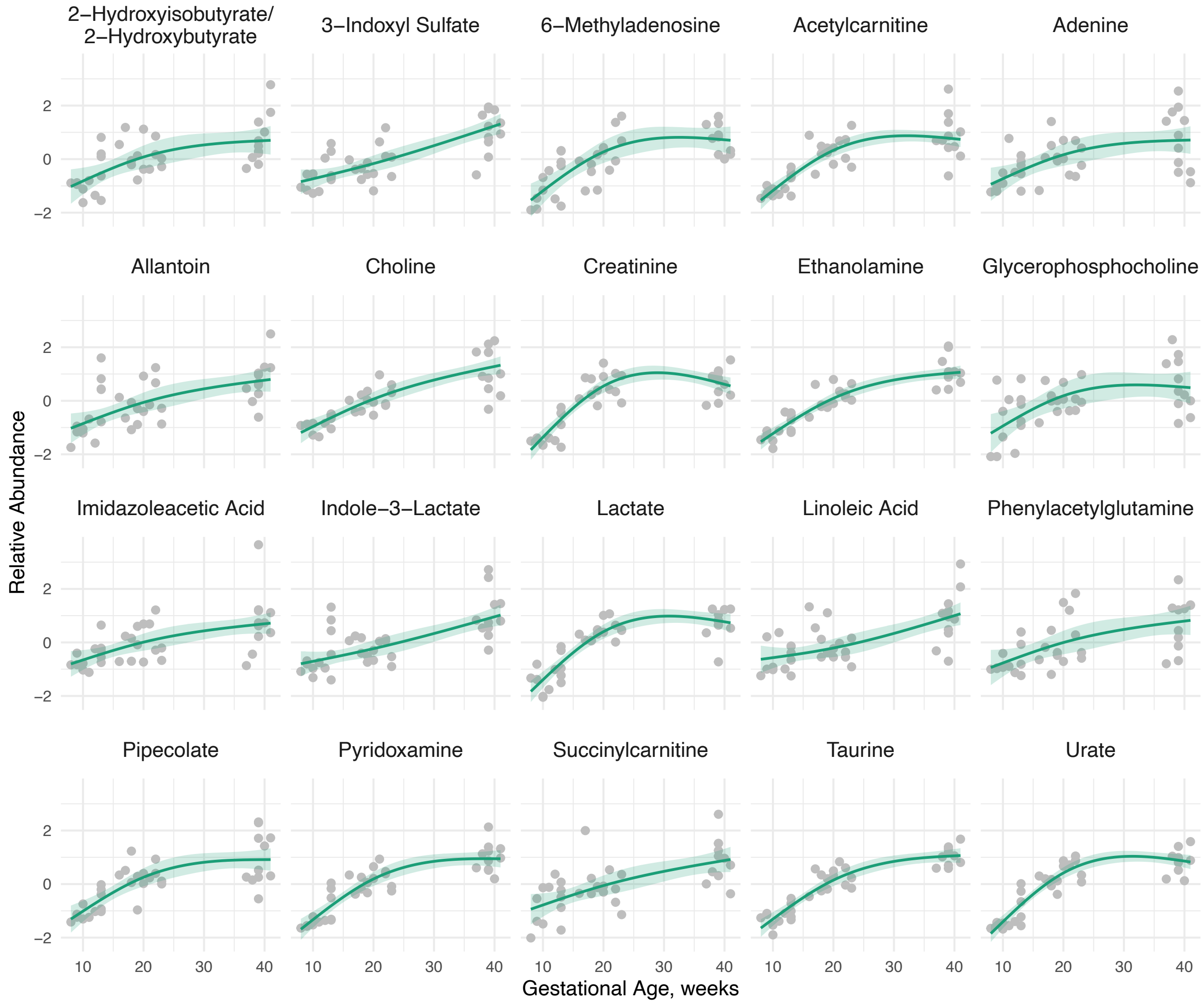


**Supplementary Figure 1. Cluster 1 metabolites with "increasing" trajectories**. Standardized metabolite relative abundances were adjusted for fetal sex and total protein. Robust models using M estimation were fit for gestational age as natural cubic spline with a single knot at the median gestational age. Estimated fits and 95% confidence intervals are presented.

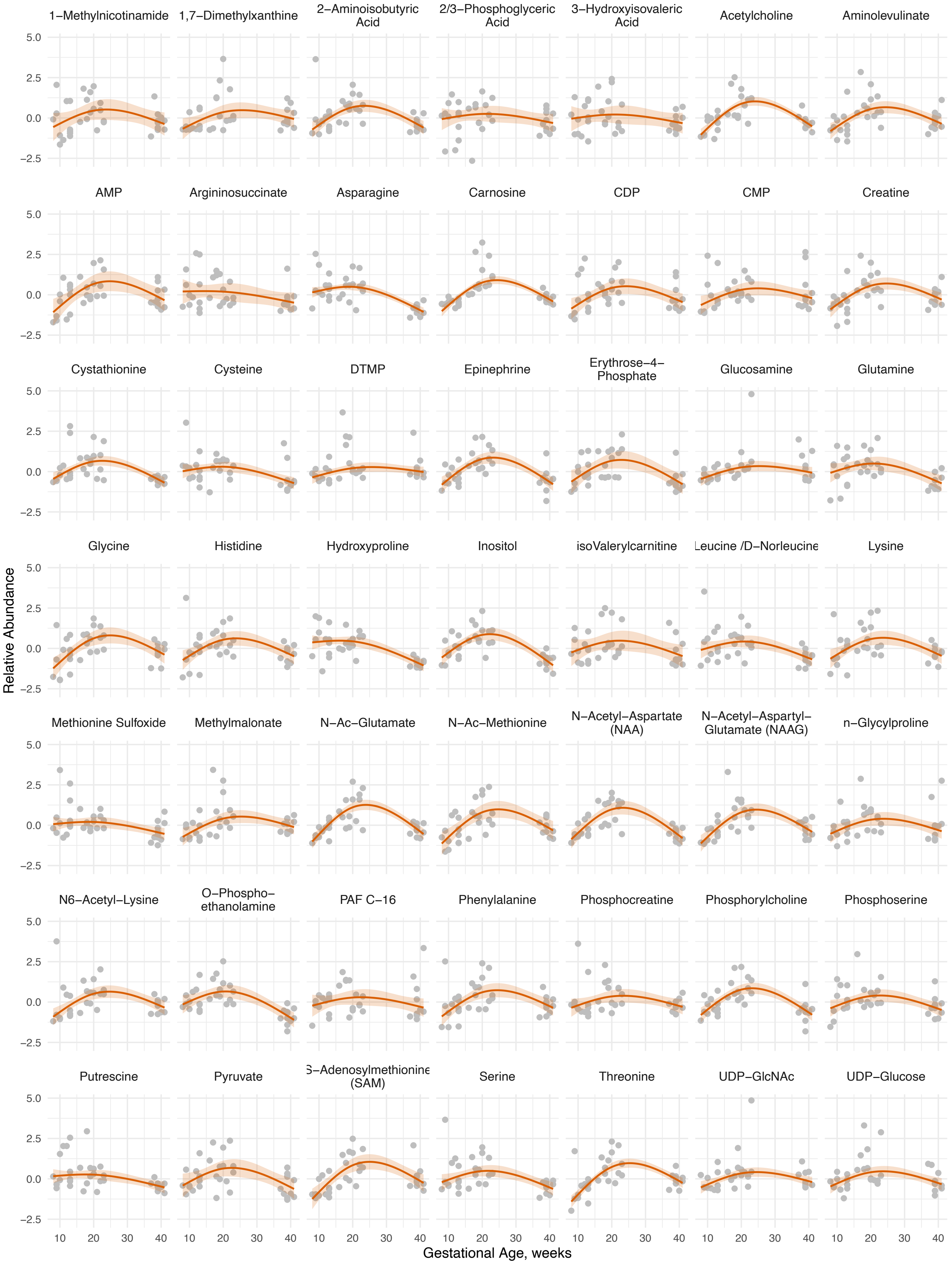
1−Methylnicotinamide
1,7−Dimethylxanthine
2−Aminoisobutyric Acid
2/3−Phosphoglyceric Acid
3−Hydroxyisovaleric Acid
Acetylcholine
Aminolevulinate
AMP
Argininosuccinate
Asparagine
Carnosine
CDP
CMP
Creatine
Cystathionine
Cysteine
DTMP
Epinephrine
Erythrose−4−Phosphate
Glucosamine
Glutamine
Glycine
Histidine
Hydroxyproline
Inositol
isoValerylcarnitine
Leucine /D−Norleucine
Lysine
Methionine Sulfoxide
Methylmalonate
N−Ac−Glutamate
N−Ac−Methionine
N−Acetyl−Aspartate (NAA)
N−Acetyl−Aspartyl−Glutamate (NAAG)
n−Glycylproline
N6−Acetyl−Lysine
O−Phospho−ethanolamine
PAF C−16
Phenylalanine
Phosphocreatine
Phosphorylcholine
Phosphoserine
Putrescine
Pyruvate
S−Adenosylmethionine (SAM)
Serine
Threonine
UDP−GlcNAc
UDP−Glucose
Relative Abundance
5.0
2.5
0.0
−2.5
10
20
30
40
Gestational Age, weeks

**Supplementary Figure 2. Cluster 2 metabolites with "decreasing into third trimester" trajectories**. Standardized metabolite relative abundances were adjusted for fetal sex and total protein. Robust models using M estimation were fit for gestational age as natural cubic spline with a single knot at the median gestational age. Estimated fits and 95% confidence intervals are presented.

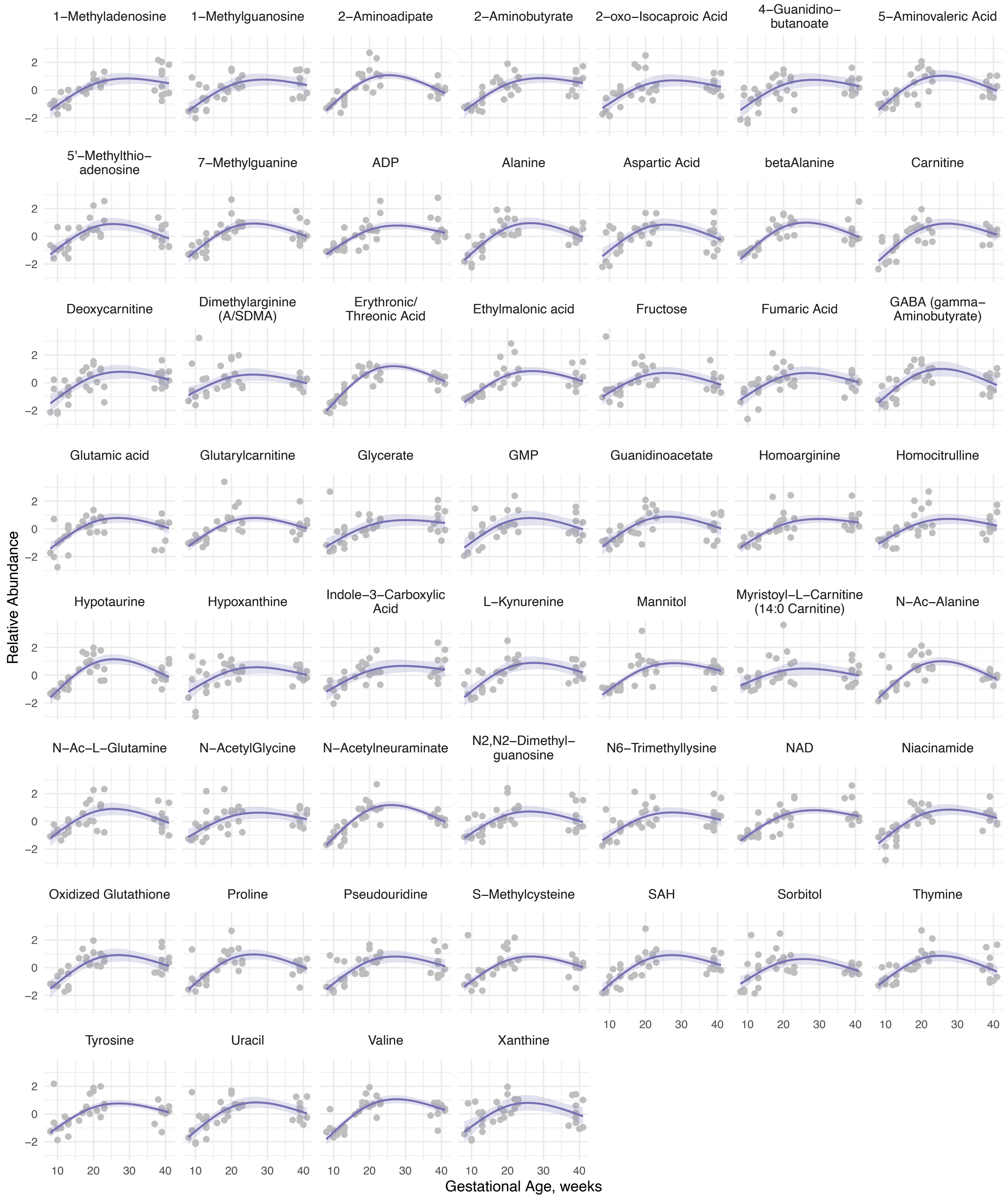


**Supplementary Figure 3. Cluster 3 metabolites with "increasing in early pregnancy" trajectories**. Standardized metabolite relative abundances were adjusted for fetal sex and total protein. Robust models using M estimation were fit for gestational age as natural cubic spline with a single knot at the median gestational age. Estimated fits and 95% confidence intervals are presented.

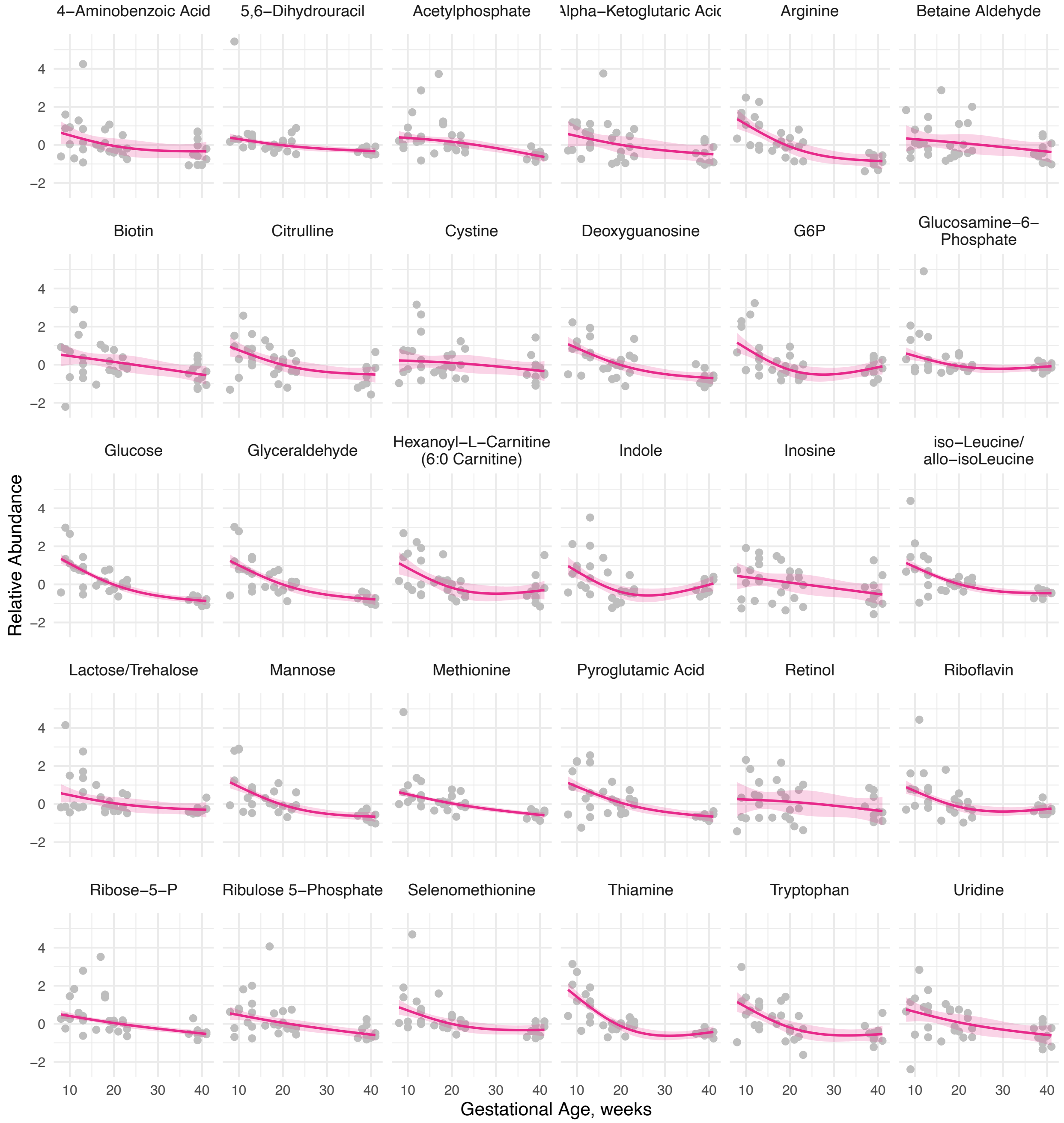


**Supplementary Figure 4. Cluster 4 metabolites with "decreasing" trajectories**. Standardized metabolite relative abundances were adjusted for fetal sex and total protein. Robust models using M estimation were fit for gestational age as natural cubic spline with a single knot at the median gestational age. Estimated fits and 95% confidence intervals are presented.

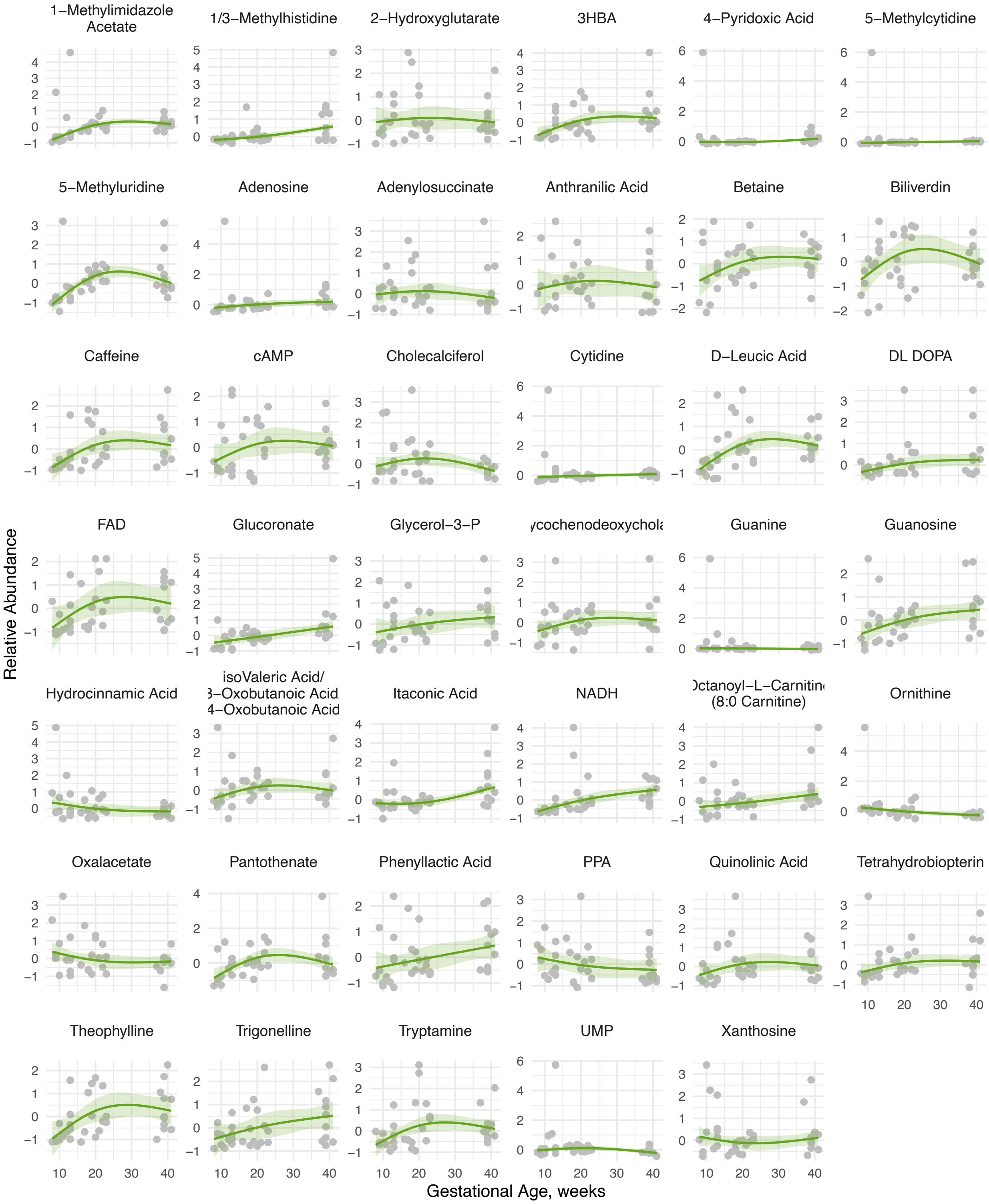


**Supplementary Figure 5. Cluster 5 metabolites with “steady” trajectories**. Standardized metabolite relative abundances were adjusted for fetal sex and total protein. Robust models using M estimation were fit for gestational age as natural cubic spline with a single knot at the median gestational age. Estimated fits and 95% confidence intervals are presented.

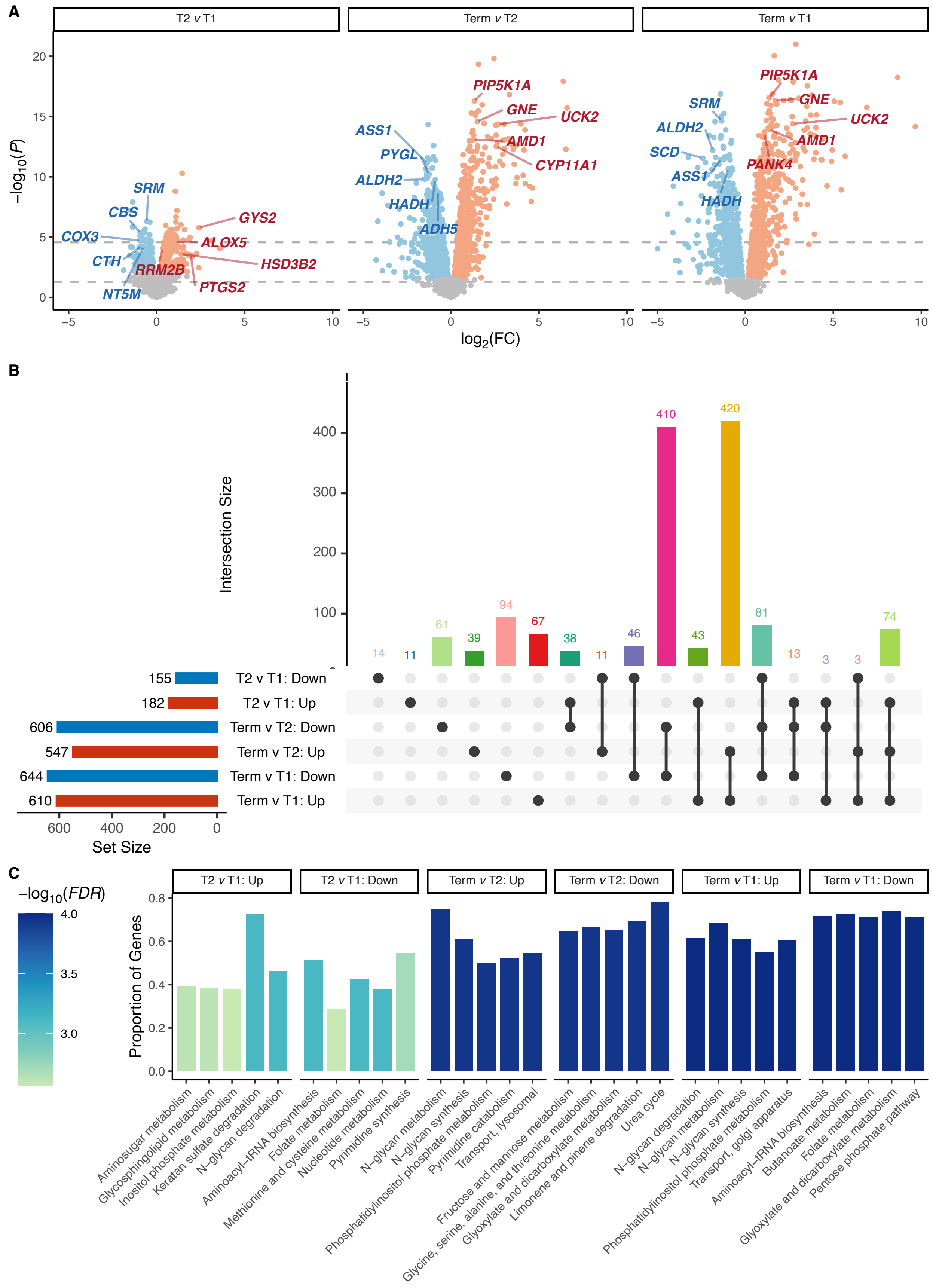


**Supplementary Figure 6**: **Transcriptomics analysis of metabolic genes**. (**A**) Volcano plot for each trimester comparison: second trimester compared to first trimester (T2 v T1), third trimester at term compared to first trimester (Term v T1), and third trimester at term compared to second trimester (Term v T2). Differentially expressed metabolic genes with FDR<0.05 are presented in color and the top 5 most upregulated and downregulated genes encoding rate limiting enzymes are labled. (**B**) An Upset plot depicting shared and distinct changes in placental metabolic gene expression. (**C**) The top 5 most upregulated and downregulated molecular subsystems were determined using rotational gene set testing. The y-axis represents the proportion of up- or downregulated genes.

**Supplementary Table 1. Differential abudance of metabolites with different trajectories in placental samples across three trimesters**. Metabolite abundances were standardized. Robust linear models using M estimation were adjusted for fetal sex and total protein to estimate the beta coefficient and standard error (SE) and corrected for false discovery rate (FDR). Comparisons with FDR<0.05 were considered significant (bold). We report here the cluster assignment for each metabolite from the metabolite trajectory clustering analysis.

| | | **Second Trimester (reference: first trimester)** | | | **Third trimester (reference: first trimester)** | | | **Third trimester (reference: second trimester)** | | |
|---|---|---|---|---|---|---|---|---|---|---|
| **Metabolite** | **Cluster** | **Beta** | **SE** | **FDR** | **Beta** | **SE** | **FDR** | **Beta** | **SE** | **FDR** |
| 1,7-Dimethylxanthine | 2 | 0.8029 | 0.3224 | **0.0428** | 0.3008 | 0.3415 | 0.4274 | -0.5021 | 0.352 | 0.3026 |
| 1-Methyladenosine | 3 | 1.3135 | 0.3051 | **0.0005** | 1.4657 | 0.3231 | **0.0004** | 0.1523 | 0.333 | 0.7518 |
| 1-Methylguanosine | 3 | 1.181 | 0.3505 | **0.0052** | 1.3054 | 0.3712 | **0.0041** | 0.1244 | 0.3826 | 0.8016 |
| 1-Methylimidazole Acetate | 5 | 0.6207 | 0.1433 | **0.0006** | 0.7539 | 0.1517 | **0.0002** | 0.1332 | 0.1564 | 0.5035 |
| 1-Methylnicotinamide | 2 | 1.0677 | 0.3971 | **0.0225** | 0.1191 | 0.4205 | 0.8147 | -0.9486 | 0.4335 | 0.0772 |
| 1/3-Methylhistidine | 5 | 0.1652 | 0.1743 | 0.3836 | 0.7805 | 0.1846 | **0.0018** | 0.6153 | 0.1903 | **0.0237** |
| 2-Aminoadipate | 3 | 1.8169 | 0.2324 | **<0.0001** | 0.7321 | 0.2462 | **0.0129** | -1.0848 | 0.2537 | **0.0018** |
| 2-Aminobutyrate | 3 | 1.5056 | 0.3469 | **0.0006** | 1.5654 | 0.3674 | **0.0006** | 0.0598 | 0.3787 | 0.9014 |
| 2-Aminoisobutyric Acid | 2 | 1.0911 | 0.2537 | **0.0006** | -0.2538 | 0.2687 | 0.4219 | -1.3449 | 0.277 | **0.0003** |
| 2-Hydroxyglutarate | 5 | 0.0796 | 0.3617 | 0.8461 | -0.0814 | 0.383 | 0.8594 | -0.161 | 0.3948 | 0.7736 |
| 2-Hydroxyisobutyrate/2-Hydroxybutyrate | 1 | 0.8457 | 0.3056 | **0.018** | 1.4578 | 0.3237 | **0.0004** | 0.6122 | 0.3336 | 0.1579 |
| 2-oxo-Isocaproic Acid | 3 | 1.1669 | 0.3627 | **0.0074** | 1.1485 | 0.3841 | **0.0128** | -0.0184 | 0.3959 | 0.9715 |
| 2/3-Phosphoglyceric Acid | 2 | 0.3113 | 0.3879 | 0.5033 | -0.3297 | 0.4108 | 0.4836 | -0.641 | 0.4235 | 0.2264 |
| 3-Hydroxyisovaleric Acid | 2 | 0.4649 | 0.4153 | 0.3386 | -0.3104 | 0.4398 | 0.5426 | -0.7753 | 0.4533 | 0.1788 |
| 3-Indoxyl Sulfate | 1 | 0.2495 | 0.2625 | 0.4136 | 1.8414 | 0.278 | **<0.0001** | 1.5919 | 0.2866 | **0.0001** |
| 3HBA | 5 | 0.4604 | 0.3207 | 0.2128 | 0.7537 | 0.3396 | 0.0599 | 0.2932 | 0.3501 | 0.5274 |
| 4-Aminobenzoic Acid | 4 | -0.4417 | 0.354 | 0.2871 | -0.7905 | 0.3749 | 0.0796 | -0.3488 | 0.3865 | 0.4902 |
| 4-Guanidinobutanoate | 3 | 1.2637 | 0.3554 | **0.0036** | 1.3111 | 0.3764 | **0.0041** | 0.0474 | 0.388 | 0.9184 |
| 4-Pyridoxic Acid | 5 | 0.0013 | 0.0444 | 0.9809 | 0.2222 | 0.047 | **0.0005** | 0.2208 | 0.0484 | **0.0008** |
| 5’-Methylthioadenosine | 3 | 1.4538 | 0.3037 | **0.0002** | 0.7475 | 0.3216 | **0.0496** | -0.7063 | 0.3315 | 0.0874 |
| 5,6-Dihydrouracil | 4 | -0.3355 | 0.1096 | **0.0115** | -0.6658 | 0.1161 | **<0.0001** | -0.3303 | 0.1197 | **0.0274** |
| 5-Aminovaleric Acid | 3 | 1.6354 | 0.2876 | **<0.0001** | 0.8582 | 0.3046 | **0.0182** | -0.7771 | 0.314 | 0.0507 |
| 5-Methylcytidine | 5 | 0.0142 | 0.0249 | 0.6321 | 0.0982 | 0.0264 | **0.0029** | 0.084 | 0.0272 | **0.0149** |
| 5-Methyluridine | 5 | 0.9564 | 0.2656 | **0.0025** | 0.7726 | 0.2813 | **0.024** | -0.1838 | 0.29 | 0.6435 |
| 6-Methyladenosine | 1 | 0.9236 | 0.3265 | **0.0161** | 1.7177 | 0.3458 | **0.0001** | 0.7941 | 0.3565 | 0.0772 |
| 7-Methylguanine | 3 | 1.6328 | 0.2738 | **<0.0001** | 1.0831 | 0.2899 | **0.0022** | -0.5497 | 0.2988 | 0.1504 |
| ADP | 3 | 1.1996 | 0.2441 | **0.0001** | 1.0779 | 0.2585 | **0.0006** | -0.1217 | 0.2665 | 0.7518 |
| AMP | 2 | 1.064 | 0.3657 | **0.0141** | 0.197 | 0.3873 | 0.6624 | -0.867 | 0.3992 | 0.0837 |
| Acetylcarnitine | 1 | 1.3296 | 0.2123 | **<0.0001** | 1.8879 | 0.2249 | **<0.0001** | 0.5584 | 0.2318 | 0.062 |
| Acetylcholine | 2 | 1.4759 | 0.2752 | **0.0001** | 0.1113 | 0.2915 | 0.742 | -1.3646 | 0.3005 | **0.0008** |
| Acetylphosphate | 4 | -0.2033 | 0.1837 | 0.352 | -1.0155 | 0.1946 | **0.0001** | -0.8122 | 0.2006 | **0.0019** |
| Adenine | 1 | 0.6216 | 0.3921 | 0.1634 | 1.3854 | 0.4152 | **0.0063** | 0.7638 | 0.428 | 0.1626 |
| Adenosine | 5 | 0.0385 | 0.16 | 0.8396 | 0.326 | 0.1694 | 0.105 | 0.2875 | 0.1746 | 0.1887 |
| Adenylosuccinate | 5 | 0.2448 | 0.353 | 0.5422 | -0.1107 | 0.3738 | 0.8054 | -0.3555 | 0.3853 | 0.4902 |

| | | Second Trimester (reference: first trimester) | | | Third trimester (reference: first trimester) | | | Third trimester (reference: second trimester) | | |
|---|---|---|---|---|---|---|---|---|---|---|
| **Metabolite** | **Cluster** | **Beta** | **SE** | **FDR** | **Beta** | **SE** | **FDR** | **Beta** | **SE** | **FDR** |
| Alanine | 3 | 1.775 | 0.3015 | **<0.0001** | 1.1026 | 0.3192 | **0.0048** | -0.6725 | 0.3291 | 0.1007 |
| Allantoin | 1 | 0.3931 | 0.4198 | 0.4244 | 1.345 | 0.4446 | **0.0123** | 0.9519 | 0.4583 | 0.0949 |
| Alpha-Ketoglutaric Acid | 4 | -0.3738 | 0.3102 | 0.3007 | -0.924 | 0.3285 | **0.0182** | -0.5501 | 0.3386 | 0.1987 |
| Aminolevulinate | 2 | 1.2576 | 0.2943 | **0.0006** | 0.2098 | 0.3117 | 0.5678 | -1.0478 | 0.3213 | **0.0135** |
| Anthranilic Acid | 5 | 0.3114 | 0.4105 | 0.5216 | 0.0552 | 0.4348 | 0.9057 | -0.2562 | 0.4482 | 0.6715 |
| Arginine | 4 | -1.0434 | 0.2516 | **0.001** | -2.0144 | 0.2664 | **<0.0001** | -0.971 | 0.2746 | **0.0062** |
| Argininosuccinate | 2 | 0.2168 | 0.3708 | 0.6168 | -0.5639 | 0.3926 | 0.217 | -0.7806 | 0.4047 | 0.118 |
| Asparagine | 2 | 0.1475 | 0.2739 | 0.6468 | -1.477 | 0.2901 | **0.0001** | -1.6245 | 0.299 | **0.0001** |
| Aspartic Acid | 3 | 1.1215 | 0.3856 | **0.0138** | 0.3849 | 0.4083 | 0.4264 | -0.7366 | 0.4209 | 0.1671 |
| Betaine | 5 | 0.5129 | 0.4298 | 0.3095 | 0.7402 | 0.4552 | 0.1665 | 0.2273 | 0.4692 | 0.7294 |
| Betaine Aldehyde | 4 | -0.2382 | 0.3376 | 0.5422 | -0.7094 | 0.3575 | 0.0858 | -0.4712 | 0.3685 | 0.3225 |
| Biliverdin | 5 | 0.4743 | 0.4393 | 0.3541 | 0.3086 | 0.4652 | 0.5678 | -0.1657 | 0.4796 | 0.8016 |
| Biotin | 4 | -0.2509 | 0.3452 | 0.5387 | -1.0428 | 0.3655 | **0.0204** | -0.7919 | 0.3768 | 0.085 |
| CDP | 2 | 0.8211 | 0.4017 | 0.0797 | 0.0584 | 0.4254 | 0.9057 | -0.7626 | 0.4384 | 0.1662 |
| CMP | 2 | 0.6343 | 0.2927 | 0.0605 | 0.1717 | 0.3099 | 0.6375 | -0.4626 | 0.3195 | 0.2543 |
| Caffeine | 5 | 0.7268 | 0.3886 | 0.109 | 0.745 | 0.4115 | 0.1183 | 0.0183 | 0.4242 | 0.9715 |
| Carnitine | 3 | 1.5894 | 0.3052 | **0.0001** | 1.3254 | 0.3232 | **0.001** | -0.2639 | 0.3332 | 0.5512 |
| Carnosine | 2 | 1.4261 | 0.1216 | **<0.0001** | 0.1479 | 0.1287 | 0.2823 | -1.2782 | 0.1327 | **<0.0001** |
| Cholecalciferol | 5 | 0.1706 | 0.3617 | 0.6889 | -0.3303 | 0.3831 | 0.4581 | -0.5009 | 0.3949 | 0.3234 |
| Choline | 1 | 0.8472 | 0.2044 | **0.0005** | 2.2651 | 0.2165 | **<0.0001** | 1.4179 | 0.2231 | **<0.0001** |
| Citrulline | 4 | -0.6277 | 0.3673 | 0.1398 | -1.2473 | 0.3889 | **0.009** | -0.6197 | 0.4009 | 0.2167 |
| Creatine | 2 | 1.3334 | 0.2922 | **0.0003** | 0.3793 | 0.3094 | 0.2931 | -0.954 | 0.3189 | **0.0208** |
| Creatinine | 1 | 1.7261 | 0.2383 | **<0.0001** | 1.8749 | 0.2523 | **<0.0001** | 0.1488 | 0.2601 | 0.6725 |
| Cystathionine | 2 | 0.8702 | 0.1598 | **0.0001** | -0.4253 | 0.1692 | **0.0298** | -1.2955 | 0.1744 | **<0.0001** |
| Cysteine | 2 | 0.3981 | 0.2244 | 0.125 | -0.7201 | 0.2376 | **0.0126** | -1.1182 | 0.2449 | **0.0006** |
| Cystine | 4 | -0.3698 | 0.3184 | 0.3298 | -0.6995 | 0.3372 | 0.0812 | -0.3296 | 0.3476 | 0.4679 |
| Cytidine | 5 | 0.0003 | 0.0957 | 0.9979 | 0.1764 | 0.1013 | 0.1491 | 0.1761 | 0.1044 | 0.1719 |
| D-Leucic Acid | 5 | 0.9682 | 0.342 | **0.0176** | 0.8475 | 0.3621 | **0.0439** | -0.1208 | 0.3733 | 0.8016 |
| DL DOPA | 5 | 0.3273 | 0.1933 | 0.1398 | 0.4617 | 0.2048 | 0.0589 | 0.1344 | 0.2111 | 0.6531 |
| DTMP | 2 | 0.5774 | 0.2495 | 0.0505 | 0.2826 | 0.2642 | 0.3459 | -0.2948 | 0.2724 | 0.4135 |
| Deoxycarnitine | 3 | 1.3161 | 0.3826 | **0.0047** | 1.2491 | 0.4052 | **0.0111** | -0.067 | 0.4177 | 0.9014 |
| Deoxyguanosine | 4 | -1.0216 | 0.2438 | **0.001** | -1.696 | 0.2582 | **<0.0001** | -0.6744 | 0.2661 | **0.04** |
| Dimethylarginine (A/SDMA) | 3 | 0.9823 | 0.3425 | **0.0161** | 0.5277 | 0.3627 | 0.2128 | -0.4546 | 0.3739 | 0.3423 |
| Epinephrine | 2 | 1.3338 | 0.2492 | **<0.0001** | -0.4116 | 0.2639 | 0.1828 | -1.7454 | 0.272 | **<0.0001** |
| Erythronic/Threonic Acid | 3 | 2.0498 | 0.2154 | **<0.0001** | 1.5437 | 0.2281 | **<0.0001** | -0.5062 | 0.2352 | 0.0808 |
| Erythrose-4-Phosphate | 2 | 0.8965 | 0.3297 | **0.0206** | -0.5185 | 0.3492 | 0.1973 | -1.415 | 0.3599 | **0.0029** |
| Ethanolamine | 1 | 1.0642 | 0.1759 | **<0.0001** | 2.2291 | 0.1862 | **<0.0001** | 1.1649 | 0.192 | **<0.0001** |
| Ethylmalonic acid | 3 | 1.4319 | 0.2563 | **<0.0001** | 1.075 | 0.2714 | **0.0011** | -0.3569 | 0.2797 | 0.3282 |
| FAD | 5 | 0.7454 | 0.4159 | 0.125 | 0.7298 | 0.4404 | 0.1534 | -0.0155 | 0.454 | 0.973 |
| Fructose | 3 | 1.2936 | 0.2789 | **0.0003** | 0.6335 | 0.2954 | 0.0722 | -0.6601 | 0.3045 | 0.0808 |

| Metabolite | Cluster | Second Trimester (reference: first trimester) | | | Third trimester (reference: first trimester) | | | Third trimester (reference: second trimester) | | |
|---|---|---|---|---|---|---|---|---|---|---|
| | | Beta | SE | FDR | Beta | SE | FDR | Beta | SE | FDR |
| Fumaric Acid | 3 | 1.3189 | 0.3385 | **0.0014** | 0.9994 | 0.3584 | **0.0204** | -0.3195 | 0.3695 | 0.5035 |
| G6P | 4 | -0.861 | 0.306 | **0.0204** | -0.7677 | 0.3241 | **0.0496** | 0.0933 | 0.3341 | 0.82 |
| GMP | 3 | 1.1135 | 0.359 | **0.0091** | 0.7602 | 0.3802 | 0.0872 | -0.3533 | 0.3919 | 0.4935 |
| Glucoronate | 5 | 0.2947 | 0.1991 | 0.1946 | 0.9504 | 0.2109 | **0.0004** | 0.6558 | 0.2173 | **0.0189** |
| Glucosamine | 2 | 0.4744 | 0.2401 | 0.0885 | 0.2036 | 0.2543 | 0.4836 | -0.2708 | 0.2621 | 0.4467 |
| Glucosamine-6-Phosphate | 4 | -0.5357 | 0.2348 | 0.0572 | -0.6442 | 0.2486 | **0.0315** | -0.1086 | 0.2563 | 0.7543 |
| Glucose | 4 | -0.9189 | 0.1724 | **0.0001** | -1.8585 | 0.1825 | **<0.0001** | -0.9396 | 0.1882 | **0.0002** |
| Glutamic acid | 3 | 1.1981 | 0.3731 | **0.0071** | 0.8406 | 0.3951 | 0.0821 | -0.3575 | 0.4073 | 0.5013 |
| Glutamine | 2 | 0.4465 | 0.3212 | 0.2352 | -0.9334 | 0.3402 | **0.024** | -1.3798 | 0.3507 | **0.0025** |
| Glutarylcarnitine | 3 | 1.4401 | 0.208 | **<0.0001** | 0.925 | 0.2203 | **0.0006** | -0.5151 | 0.2271 | 0.0796 |
| Glyceraldehyde | 4 | -0.8951 | 0.2043 | **0.0008** | -1.7149 | 0.2163 | **<0.0001** | -0.8197 | 0.223 | **0.0032** |
| Glycerate | 3 | 0.9934 | 0.3098 | **0.0073** | 1.3025 | 0.328 | **0.0015** | 0.3091 | 0.3381 | 0.4902 |
| Glycerol-3-P | 5 | 0.0956 | 0.3786 | 0.8367 | 0.6575 | 0.4009 | 0.1611 | 0.5619 | 0.4133 | 0.2904 |
| Glycerophosphocholine | 1 | 0.727 | 0.4247 | 0.1398 | 1.3062 | 0.4498 | **0.0164** | 0.5793 | 0.4636 | 0.3282 |
| Glycine | 2 | 1.129 | 0.3485 | **0.0078** | 0.1308 | 0.3691 | 0.7786 | -0.9983 | 0.3805 | **0.033** |
| Glycochenodeoxycholate | 5 | 0.2716 | 0.3376 | 0.5003 | 0.3411 | 0.3576 | 0.4184 | 0.0695 | 0.3686 | 0.8934 |
| Guanidinoacetate | 3 | 1.3624 | 0.333 | **0.0009** | 0.7502 | 0.3526 | 0.0722 | -0.6123 | 0.3635 | 0.1842 |
| Guanine | 5 | -0.0084 | 0.0301 | 0.8213 | -0.0371 | 0.0318 | 0.3472 | -0.0286 | 0.0328 | 0.4935 |
| Guanosine | 5 | 0.1684 | 0.3257 | 0.6504 | 0.7727 | 0.3449 | 0.063 | 0.6043 | 0.3555 | 0.1762 |
| Hexanoyl-L-Carnitine (6:0 Carnitine) | 4 | -0.8275 | 0.3766 | 0.0605 | -1.0971 | 0.3989 | **0.0218** | -0.2696 | 0.4111 | 0.6319 |
| Histidine | 2 | 0.6902 | 0.2935 | 0.0503 | -0.317 | 0.3108 | 0.4 | -1.0072 | 0.3204 | **0.0138** |
| Homoarginine | 3 | 1.2504 | 0.2227 | **<0.0001** | 1.4307 | 0.2359 | **<0.0001** | 0.1803 | 0.2431 | 0.5784 |
| Homocitrulline | 3 | 1.2894 | 0.3224 | **0.0011** | 0.9472 | 0.3415 | **0.0202** | -0.3422 | 0.352 | 0.4772 |
| Hydrocinnamic Acid | 5 | -0.1741 | 0.2327 | 0.5292 | -0.4213 | 0.2464 | 0.1478 | -0.2472 | 0.254 | 0.4598 |
| Hydroxyproline | 2 | 0.0181 | 0.2589 | 0.9559 | -1.5635 | 0.2742 | **<0.0001** | -1.5816 | 0.2826 | **<0.0001** |
| Hypotaurine | 3 | 1.9198 | 0.2956 | **<0.0001** | 0.9551 | 0.313 | **0.0123** | -0.9647 | 0.3227 | **0.0216** |
| Hypoxanthine | 3 | 1.1391 | 0.3278 | **0.0047** | 0.8672 | 0.3471 | **0.0419** | -0.2719 | 0.3578 | 0.5579 |
| Imidazoleacetic Acid | 1 | 0.5052 | 0.2591 | 0.0889 | 1.309 | 0.2744 | **0.0002** | 0.8038 | 0.2828 | **0.0274** |
| Indole | 4 | -1.0733 | 0.3188 | **0.0069** | -0.6527 | 0.3376 | 0.0943 | 0.4206 | 0.348 | 0.3347 |
| Indole-3-Carboxylic Acid | 3 | 1.1075 | 0.3316 | **0.0052** | 1.2267 | 0.3512 | **0.0041** | 0.1192 | 0.362 | 0.8016 |
| Indole-3-Lactate | 1 | 0.3574 | 0.2499 | 0.2128 | 1.6887 | 0.2646 | **<0.0001** | 1.3313 | 0.2728 | **0.0004** |
| Inosine | 4 | -0.4646 | 0.4568 | 0.3836 | -1.0019 | 0.4838 | 0.081 | -0.5373 | 0.4987 | 0.408 |
| Inositol | 2 | 1.001 | 0.2505 | **0.0012** | -0.9398 | 0.2652 | **0.0041** | -1.9408 | 0.2734 | **<0.0001** |
| Itaconic Acid | 5 | 0.0276 | 0.135 | 0.8461 | 0.8889 | 0.143 | **<0.0001** | 0.8612 | 0.1474 | **0.0001** |
| L-Kynurenine | 3 | 1.4759 | 0.3153 | **0.0003** | 1.2681 | 0.3339 | **0.0019** | -0.2078 | 0.3442 | 0.6617 |
| Lactate | 1 | 1.5805 | 0.1894 | **<0.0001** | 2.1346 | 0.2005 | **<0.0001** | 0.5541 | 0.2067 | **0.0371** |
| Lactose/Trehalose | 4 | -0.4656 | 0.2482 | 0.1157 | -0.8584 | 0.2629 | **0.0079** | -0.3928 | 0.271 | 0.2435 |
| Leucine /D-Norleucine | 2 | 0.344 | 0.3809 | 0.453 | -0.8686 | 0.4034 | 0.0789 | -1.2126 | 0.4158 | **0.0216** |
| Linoleic Acid | 1 | 0.472 | 0.2827 | 0.1398 | 1.6815 | 0.2994 | **<0.0001** | 1.2095 | 0.3086 | **0.003** |
| Lysine | 2 | 0.9051 | 0.3552 | **0.0306** | -0.2306 | 0.3762 | 0.6013 | -1.1357 | 0.3878 | **0.0218** |

| | | Second Trimester (reference: first trimester) | | | Third trimester (reference: first trimester) | | | Third trimester (reference: second trimester) | | |
|---|---|---|---|---|---|---|---|---|---|---|
| **Metabolite** | **Cluster** | **Beta** | **SE** | **FDR** | **Beta** | **SE** | **FDR** | **Beta** | **SE** | **FDR** |
| Mannitol | 3 | 1.5936 | 0.1505 | **<0.0001** | 1.4107 | 0.1594 | **<0.0001** | -0.1829 | 0.1643 | 0.4054 |
| Mannose | 4 | -0.6521 | 0.2315 | **0.0199** | -1.498 | 0.2451 | **<0.0001** | -0.8459 | 0.2527 | **0.0085** |
| Methionine | 4 | -0.4425 | 0.1696 | **0.0284** | -1.1852 | 0.1797 | **<0.0001** | -0.7428 | 0.1852 | **0.0021** |
| Methionine Sulfoxide | 2 | 0.0928 | 0.292 | 0.7999 | -0.724 | 0.3093 | 0.0602 | -0.8168 | 0.3188 | **0.04** |
| Methylmalonate | 2 | 1.0269 | 0.2631 | **0.0019** | 0.3473 | 0.2786 | 0.2659 | -0.6796 | 0.2872 | 0.0669 |
| Myristoyl-L-Carnitine (14:0 Carnitine) | 3 | 0.9771 | 0.3331 | **0.0141** | 0.5002 | 0.3528 | 0.2163 | -0.4769 | 0.3637 | 0.3233 |
| N-Ac-Alanine | 3 | 1.7539 | 0.2837 | **<0.0001** | 0.6292 | 0.3004 | 0.0812 | -1.1246 | 0.3096 | **0.005** |
| N-Ac-Glutamate | 2 | 1.3974 | 0.2804 | **0.0001** | -0.0655 | 0.297 | 0.8539 | -1.4629 | 0.3061 | **0.0004** |
| N-Ac-L-Glutamine | 3 | 1.2558 | 0.3886 | **0.0071** | 0.6916 | 0.4115 | 0.1491 | -0.5642 | 0.4242 | 0.3164 |
| N-Ac-Methionine | 2 | 1.4105 | 0.3527 | **0.0012** | 0.3246 | 0.3735 | 0.4546 | -1.0859 | 0.385 | **0.0274** |
| N-Acetyl-Aspartate (NAA) | 2 | 1.3544 | 0.2531 | **0.0001** | -0.3622 | 0.268 | 0.2389 | -1.7165 | 0.2763 | **<0.0001** |
| N-Acetyl-Aspartyl-Glutamate (NAAG) | 2 | 1.5541 | 0.2302 | **<0.0001** | 0.3673 | 0.2438 | 0.1828 | -1.1868 | 0.2513 | **0.0004** |
| N-AcetylGlycine | 3 | 1.0361 | 0.3154 | **0.0072** | 0.8909 | 0.334 | **0.0234** | -0.1452 | 0.3443 | 0.759 |
| N-Acetylneuraminate | 3 | 1.9602 | 0.2387 | **<0.0001** | 1.2246 | 0.2528 | **0.0001** | -0.7356 | 0.2606 | **0.0274** |
| N2,N2-Dimethylguanosine | 3 | 1.2527 | 0.3706 | **0.0052** | 0.7666 | 0.3925 | 0.0927 | -0.4862 | 0.4046 | 0.359 |
| N6-Acetyl-Lysine | 2 | 1.0294 | 0.3055 | **0.006** | 0.1934 | 0.3236 | 0.607 | -0.836 | 0.3335 | **0.0444** |
| N6-Trimethyllysine | 3 | 1.2007 | 0.3569 | **0.0052** | 1.0529 | 0.378 | **0.0202** | -0.1478 | 0.3896 | 0.7847 |
| NAD | 3 | 1.2363 | 0.2306 | **<0.0001** | 1.2829 | 0.2442 | **<0.0001** | 0.0465 | 0.2517 | 0.8934 |
| NADH | 5 | 0.4269 | 0.2271 | 0.1076 | 1.0321 | 0.2405 | **0.0005** | 0.6052 | 0.248 | 0.0577 |
| Niacinamide | 3 | 1.6355 | 0.2713 | **<0.0001** | 1.4794 | 0.2873 | **0.0001** | -0.1561 | 0.2962 | 0.7065 |
| Octanoyl-L-Carnitine (8:0 Carnitine) | 5 | 0.1762 | 0.2391 | 0.5282 | 0.728 | 0.2532 | **0.0195** | 0.5518 | 0.261 | 0.0992 |
| Ornithine | 5 | -0.2144 | 0.1056 | 0.0889 | -0.4809 | 0.1118 | **0.0005** | -0.2665 | 0.1153 | 0.061 |
| Oxalacetate | 5 | -0.0999 | 0.4137 | 0.8396 | -0.3961 | 0.4382 | 0.4423 | -0.2962 | 0.4516 | 0.6272 |
| Oxidized Glutathione | 3 | 1.8154 | 0.2422 | **<0.0001** | 1.1799 | 0.2565 | **0.0003** | -0.6354 | 0.2644 | 0.0605 |
| PAF C-16 | 2 | 0.5634 | 0.3789 | 0.1925 | -0.0653 | 0.4013 | 0.895 | -0.6287 | 0.4137 | 0.2336 |
| PPA | 5 | -0.0891 | 0.3327 | 0.8305 | -0.4521 | 0.3524 | 0.2714 | -0.3629 | 0.3632 | 0.47 |
| Pantothenate | 5 | 0.5724 | 0.3782 | 0.1844 | 0.4316 | 0.4005 | 0.3577 | -0.1408 | 0.4129 | 0.8016 |
| Phenylacetylglutamine | 1 | 0.5204 | 0.3529 | 0.1992 | 1.4985 | 0.3737 | **0.0011** | 0.978 | 0.3852 | **0.0481** |
| Phenylalanine | 2 | 1.0796 | 0.3677 | **0.0141** | 0.0495 | 0.3894 | 0.9057 | -1.0301 | 0.4013 | **0.0432** |
| Phenyllactic Acid | 5 | 0.2257 | 0.3861 | 0.6168 | 0.8255 | 0.4089 | 0.0842 | 0.5998 | 0.4214 | 0.2672 |
| Phosphocreatine | 2 | 0.7714 | 0.2675 | **0.0155** | -0.1267 | 0.2833 | 0.7149 | -0.898 | 0.292 | **0.0226** |
| Phosphorylcholine | 2 | 1.3225 | 0.24 | **<0.0001** | -0.4325 | 0.2541 | 0.1491 | -1.755 | 0.262 | **<0.0001** |
| Phosphoserine | 2 | 0.3736 | 0.2602 | 0.2151 | -0.3899 | 0.2755 | 0.2215 | -0.7635 | 0.284 | **0.0299** |
| Pipecolate | 1 | 1.1259 | 0.2629 | **0.0005** | 1.8667 | 0.2784 | **<0.0001** | 0.7407 | 0.287 | **0.0467** |
| Proline | 3 | 1.7305 | 0.2257 | **<0.0001** | 0.9532 | 0.2391 | **0.0015** | -0.7773 | 0.2464 | **0.0183** |
| Pseudouridine | 3 | 1.3115 | 0.3746 | **0.0039** | 1.1353 | 0.3967 | **0.0181** | -0.1761 | 0.4089 | 0.7543 |
| Putrescine | 2 | 0.0448 | 0.2586 | 0.8835 | -0.701 | 0.2739 | **0.0309** | -0.7458 | 0.2823 | **0.033** |
| Pyridoxamine | 1 | 1.3085 | 0.2156 | **<0.0001** | 2.2207 | 0.2283 | **<0.0001** | 0.9123 | 0.2354 | **0.0032** |
| Pyroglutamic Acid | 4 | -0.6925 | 0.2997 | 0.0575 | -1.7341 | 0.3174 | **0.0001** | -1.0416 | 0.3271 | **0.0123** |
| Pyruvate | 2 | 0.7766 | 0.3671 | 0.0704 | -0.4502 | 0.3887 | 0.3127 | -1.2268 | 0.4007 | **0.0185** |

| | | Second Trimester (reference: first trimester) | | | Third trimester (reference: first trimester) | | | Third trimester (reference: second trimester) | | |
|---|---|---|---|---|---|---|---|---|---|---|
| **Metabolite** | **Cluster** | **Beta** | **SE** | **FDR** | **Beta** | **SE** | **FDR** | **Beta** | **SE** | **FDR** |
| Quinolinic Acid | 5 | 0.5731 | 0.3029 | 0.1056 | 0.4763 | 0.3207 | 0.1921 | -0.0968 | 0.3306 | 0.8161 |
| Retinol | 4 | -0.2777 | 0.4154 | 0.564 | -0.6615 | 0.4399 | 0.1941 | -0.3838 | 0.4534 | 0.516 |
| Riboflavin | 4 | -0.7278 | 0.2563 | **0.0178** | -0.8451 | 0.2714 | **0.0111** | -0.1173 | 0.2797 | 0.7543 |
| Ribose-5-P | 4 | -0.3857 | 0.1631 | 0.0503 | -0.9451 | 0.1727 | **<0.0001** | -0.5594 | 0.178 | **0.0125** |
| Ribulose 5-Phosphate | 4 | -0.3391 | 0.2479 | 0.2497 | -1.0563 | 0.2625 | **0.0013** | -0.7172 | 0.2706 | **0.0312** |
| S-Adenosylmethionine (SAM) | 2 | 1.6364 | 0.3048 | **<0.0001** | 0.5034 | 0.3228 | 0.1792 | -1.1331 | 0.3327 | **0.0097** |
| S-Methylcysteine | 3 | 1.4643 | 0.298 | **0.0001** | 0.8383 | 0.3156 | **0.0263** | -0.6261 | 0.3253 | 0.1265 |
| SAH | 3 | 1.47 | 0.3087 | **0.0002** | 1.2298 | 0.327 | **0.0021** | -0.2402 | 0.337 | 0.6021 |
| Selenomethionine | 4 | -0.525 | 0.2092 | **0.0344** | -0.9183 | 0.2215 | **0.001** | -0.3933 | 0.2283 | 0.1633 |
| Serine | 2 | 0.6549 | 0.2992 | 0.0629 | -0.6473 | 0.3169 | 0.0836 | -1.3022 | 0.3266 | **0.0024** |
| Sorbitol | 3 | 1.3123 | 0.3064 | **0.0007** | 0.6571 | 0.3245 | 0.0842 | -0.6551 | 0.3344 | 0.1101 |
| Succinylcarnitine | 1 | 0.4994 | 0.2967 | 0.1398 | 1.8122 | 0.3143 | **<0.0001** | 1.3128 | 0.3239 | **0.0026** |
| Taurine | 1 | 1.2926 | 0.1368 | **<0.0001** | 2.3715 | 0.1448 | **<0.0001** | 1.0788 | 0.1493 | **<0.0001** |
| Tetrahydrobiopterin | 5 | 0.332 | 0.2405 | 0.2151 | 0.4273 | 0.2547 | 0.1681 | 0.0953 | 0.2625 | 0.7985 |
| Theophylline | 5 | 0.8087 | 0.4173 | 0.0958 | 0.8644 | 0.4419 | 0.0939 | 0.0558 | 0.4555 | 0.9184 |
| Thiamine | 4 | -1.3872 | 0.1305 | **<0.0001** | -1.8215 | 0.1382 | **<0.0001** | -0.4344 | 0.1424 | **0.0096** |
| Threonine | 2 | 1.5331 | 0.3067 | **0.0001** | 0.5561 | 0.3248 | 0.1472 | -0.977 | 0.3348 | **0.0233** |
| Thymine | 3 | 1.317 | 0.3194 | **0.0008** | 0.5242 | 0.3382 | 0.1799 | -0.7928 | 0.3486 | 0.0748 |
| Trigonelline | 5 | 0.286 | 0.3886 | 0.5282 | 0.845 | 0.4115 | 0.0812 | 0.5589 | 0.4242 | 0.3195 |
| Tryptamine | 5 | 0.6096 | 0.368 | 0.1608 | 0.4846 | 0.3897 | 0.276 | -0.125 | 0.4017 | 0.8161 |
| Tryptophan | 4 | -0.8166 | 0.2766 | **0.0141** | -1.3286 | 0.2929 | **0.0004** | -0.512 | 0.3019 | 0.1788 |
| Tyrosine | 3 | 1.3707 | 0.3213 | **0.0007** | 0.9484 | 0.3403 | **0.0203** | -0.4223 | 0.3508 | 0.3632 |
| UDP-GlcNAc | 2 | 0.6554 | 0.2225 | **0.0142** | 0.1507 | 0.2356 | 0.5695 | -0.5048 | 0.2429 | 0.0992 |
| UDP-Glucose | 2 | 0.8999 | 0.2822 | **0.0083** | 0.0432 | 0.2989 | 0.902 | -0.8568 | 0.3081 | **0.033** |
| UMP | 5 | 0.0835 | 0.1039 | 0.5033 | -0.208 | 0.11 | 0.116 | -0.2915 | 0.1134 | **0.0372** |
| Uracil | 3 | 1.4612 | 0.322 | **0.0004** | 1.2039 | 0.341 | **0.0041** | -0.2573 | 0.3515 | 0.5784 |
| Urate | 1 | 1.6303 | 0.2198 | **<0.0001** | 2.1853 | 0.2327 | **<0.0001** | 0.555 | 0.2399 | 0.0669 |
| Uridine | 4 | -0.5827 | 0.3292 | 0.1304 | -1.3367 | 0.3486 | **0.0023** | -0.7539 | 0.3594 | 0.0874 |
| Valine | 3 | 1.8776 | 0.171 | **<0.0001** | 1.6537 | 0.1811 | **<0.0001** | -0.2238 | 0.1867 | 0.3657 |
| Xanthine | 3 | 1.173 | 0.3784 | **0.0091** | 0.5771 | 0.4008 | 0.2128 | -0.5959 | 0.4131 | 0.2626 |
| Xanthosine | 5 | -0.4467 | 0.2354 | 0.1051 | -0.12 | 0.2493 | 0.6924 | 0.3267 | 0.257 | 0.3193 |
| betaAlanine | 3 | 1.8459 | 0.2276 | **<0.0001** | 1.0623 | 0.241 | **0.0005** | -0.7836 | 0.2485 | **0.0187** |
| cAMP | 5 | 0.4636 | 0.4336 | 0.3631 | 0.3851 | 0.4592 | 0.4651 | -0.0785 | 0.4733 | 0.9003 |
| gamma-Aminobutyrate | 3 | 1.7906 | 0.2867 | **<0.0001** | 0.8274 | 0.3036 | **0.0218** | -0.9632 | 0.313 | **0.0183** |
| iso-Leucine /allo-isoLeucine | 4 | -0.835 | 0.1616 | **0.0001** | -1.4052 | 0.1712 | **<0.0001** | -0.5701 | 0.1765 | **0.0097** |
| isoValeric Acid /3-Oxobutanoic Acid/4-Oxobutanoic Acid | 5 | 0.575 | 0.2713 | 0.0733 | 0.312 | 0.2873 | 0.3528 | -0.2631 | 0.2962 | 0.4935 |
| isoValerylcarnitine | 2 | 0.7175 | 0.4314 | 0.1475 | -0.373 | 0.4569 | 0.4779 | -1.0905 | 0.471 | 0.0669 |
| n-Glycylproline | 2 | 0.6826 | 0.3173 | 0.0605 | -0.0073 | 0.3361 | 0.9829 | -0.6899 | 0.3464 | 0.1078 |
| o-Phosphoethanolamine | 2 | 0.8152 | 0.2406 | **0.0052** | -1.1569 | 0.2548 | **0.0004** | -1.9721 | 0.2627 | **<0.0001** |

**Supplementary Table 2**. Directional Goeman's global tests (GGT) of Recon3D molecular subsystems (FDR<0.05). Metabolites in these subsystems are consistently up- or downregulated in the same direction. Models were adjusted for fetal sex and total protein.

| Contrast | Direction | Molecular Subsystem | Fold Enrichment | FDR |
|---|---|---|---|---|
| **T2vT1** | Up | Alanine and aspartate metabolism | 5.4 | 2.87E-03 |
| **T2vT1** | Up | Aminoacyl-tRNA biosynthesis | 4.9 | 1.25E-02 |
| **T2vT1** | Up | Aminosugar metabolism | 7.6 | 1.28E-04 |
| **T2vT1** | Up | Arachidonic acid metabolism | 10.2 | 7.35E-06 |
| **T2vT1** | Up | Arginine and proline metabolism | 8.3 | 3.19E-05 |
| **T2vT1** | Up | Beta-Alanine metabolism | 12.6 | 1.65E-07 |
| **T2vT1** | Up | Bile acid synthesis | 8.5 | 2.42E-04 |
| **T2vT1** | Up | Butanoate metabolism | 5.8 | 1.64E-03 |
| **T2vT1** | Up | Cholesterol metabolism | 9.4 | 4.43E-05 |
| **T2vT1** | Up | Citric acid cycle | 10.5 | 1.24E-06 |
| **T2vT1** | Up | CoA synthesis | 6.4 | 4.96E-04 |
| **T2vT1** | Up | Fatty acid oxidation | 10.8 | 6.55E-07 |
| **T2vT1** | Up | Fructose and mannose metabolism | 4.7 | 3.91E-03 |
| **T2vT1** | Up | Glutamate metabolism | 13.1 | 1.65E-07 |
| **T2vT1** | Up | Glutathione metabolism | 11.6 | 4.34E-07 |
| **T2vT1** | Up | Glycerophospholipid metabolism | 11.3 | 8.61E-07 |
| **T2vT1** | Up | Glycine, serine, alanine, and threonine metabolism | 8.1 | 4.79E-05 |
| **T2vT1** | Up | Glycolysis/gluconeogenesis | 6.1 | 8.91E-04 |
| **T2vT1** | Up | Heme synthesis | 8.7 | 2.59E-05 |
| **T2vT1** | Up | Histidine metabolism | 11.2 | 1.32E-06 |
| **T2vT1** | Up | Lysine metabolism | 13.0 | 1.65E-07 |
| **T2vT1** | Up | Methionine and cysteine metabolism | 5.8 | 1.91E-03 |
| **T2vT1** | Up | NAD metabolism | 12.9 | 1.65E-07 |
| **T2vT1** | Up | Nucleotide interconversion | 6.6 | 1.45E-03 |
| **T2vT1** | Up | Nucleotide metabolism | 7.0 | 3.32E-05 |
| **T2vT1** | Up | Nucleotide salvage pathway | 5.2 | 4.30E-03 |
| **T2vT1** | Up | Pentose phosphate pathway | 6.0 | 1.85E-03 |
| **T2vT1** | Up | Phenylalanine metabolism | 10.7 | 2.59E-06 |
| **T2vT1** | Up | Propanoate metabolism | 8.3 | 3.25E-04 |
| **T2vT1** | Up | Protein degradation | 5.1 | 1.08E-02 |
| **T2vT1** | Up | Purine catabolism | 5.9 | 3.49E-03 |
| **T2vT1** | Up | Pyrimidine catabolism | 5.8 | 2.11E-04 |
| **T2vT1** | Up | Pyrimidine synthesis | 6.9 | 1.81E-04 |
| **T2vT1** | Up | Pyruvate metabolism | 10.2 | 8.82E-07 |
| **T2vT1** | Up | Selenoamino acid metabolism | 5.5 | 2.30E-03 |
| **T2vT1** | Up | Sphingolipid metabolism | 9.2 | 9.82E-07 |
| **T2vT1** | Up | Starch and sucrose metabolism | 4.1 | 3.89E-03 |

| Contrast | Direction | Molecular Subsystem | Fold Enrichment | FDR |
|---|---|---|---|---|
| **T2vT1** | Up | Transport, endoplasmic reticular | 4.8 | 1.97E-03 |
| **T2vT1** | Up | Transport, extracellular | 8.1 | 3.25E-04 |
| **T2vT1** | Up | Transport, golgi apparatus | 5.2 | 2.30E-03 |
| **T2vT1** | Up | Transport, lysosomal | 7.5 | 4.46E-04 |
| **T2vT1** | Up | Transport, mitochondrial | 5.0 | 5.92E-03 |
| **T2vT1** | Up | Transport, peroxisomal | 13.1 | 8.11E-10 |
| **T2vT1** | Up | Tryptophan metabolism | 10.6 | 1.77E-06 |
| **T2vT1** | Up | Tyrosine metabolism | 12.6 | 1.71E-07 |
| **T2vT1** | Up | Urea cycle | 11.9 | 1.65E-07 |
| **T2vT1** | Up | Valine, leucine, and isoleucine metabolism | 10.8 | 8.61E-07 |
| **T2vT1** | Up | Xenobiotics metabolism | 8.4 | 2.73E-04 |
| **TermvT1** | Up | Aminosugar metabolism | 6.3 | 4.89E-04 |
| **TermvT1** | Up | Arachidonic acid metabolism | 8.2 | 1.37E-04 |
| **TermvT1** | Up | Arginine and proline metabolism | 3.2 | 2.38E-02 |
| **TermvT1** | Up | Beta-Alanine metabolism | 5.9 | 3.11E-03 |
| **TermvT1** | Up | Bile acid synthesis | 7.6 | 3.18E-04 |
| **TermvT1** | Up | Butanoate metabolism | 7.3 | 2.12E-04 |
| **TermvT1** | Up | Cholesterol metabolism | 9.9 | 1.01E-04 |
| **TermvT1** | Up | Citric acid cycle | 6.7 | 3.20E-04 |
| **TermvT1** | Up | Fatty acid oxidation | 9.4 | 1.01E-04 |
| **TermvT1** | Up | Fructose and mannose metabolism | 5.7 | 1.01E-04 |
| **TermvT1** | Up | Glutamate metabolism | 6.4 | 1.00E-03 |
| **TermvT1** | Up | Glutathione metabolism | 7.0 | 4.24E-04 |
| **TermvT1** | Up | Glycerophospholipid metabolism | 6.7 | 4.46E-04 |
| **TermvT1** | Up | Glycine, serine, alanine, and threonine metabolism | 3.8 | 2.53E-02 |
| **TermvT1** | Up | Glycolysis/gluconeogenesis | 4.7 | 5.11E-03 |
| **TermvT1** | Up | Heme synthesis | 8.6 | 1.11E-04 |
| **TermvT1** | Up | Histidine metabolism | 7.9 | 3.20E-04 |
| **TermvT1** | Up | Lysine metabolism | 9.0 | 1.37E-04 |
| **TermvT1** | Up | N-glycan synthesis | 6.8 | 2.69E-04 |
| **TermvT1** | Up | NAD metabolism | 8.5 | 1.38E-04 |
| **TermvT1** | Up | Pentose phosphate pathway | 3.5 | 1.90E-02 |
| **TermvT1** | Up | Phenylalanine metabolism | 7.3 | 4.24E-04 |
| **TermvT1** | Up | Propanoate metabolism | 8.7 | 1.37E-04 |
| **TermvT1** | Up | Purine catabolism | 4.6 | 1.55E-02 |
| **TermvT1** | Up | Pyruvate metabolism | 9.0 | 1.37E-04 |
| **TermvT1** | Up | Sphingolipid metabolism | 3.4 | 2.53E-02 |
| **TermvT1** | Up | Starch and sucrose metabolism | 5.2 | 2.30E-04 |
| **TermvT1** | Up | Transport, endoplasmic reticular | 4.4 | 2.79E-03 |
| **TermvT1** | Up | Transport, peroxisomal | 10.8 | 2.95E-05 |

| Contrast | Direction | Molecular Subsystem | Fold Enrichment | FDR |
|---|---|---|---|---|
| **TermvT1** | Up | Tryptophan metabolism | 7.5 | 3.20E-04 |
| **TermvT1** | Up | Tyrosine metabolism | 7.5 | 2.94E-04 |
| **TermvT1** | Up | Urea cycle | 4.5 | 8.02E-03 |
| **TermvT1** | Up | Valine, leucine, and isoleucine metabolism | 8.3 | 1.37E-04 |
| **TermvT1** | Up | Xenobiotics metabolism | 8.6 | 1.37E-04 |
| **TermvT2** | Down | Alanine and aspartate metabolism | 8.1 | 8.44E-04 |
| **TermvT2** | Down | Aminoacyl-tRNA biosynthesis | 8.5 | 8.44E-04 |
| **TermvT2** | Down | Aminosugar metabolism | 3.8 | 1.40E-02 |
| **TermvT2** | Down | Arginine and proline metabolism | 7.3 | 1.02E-03 |
| **TermvT2** | Down | Beta-Alanine metabolism | 6.5 | 1.02E-03 |
| **TermvT2** | Down | Citric acid cycle | 3.6 | 4.46E-02 |
| **TermvT2** | Down | Glutamate metabolism | 6.3 | 2.84E-03 |
| **TermvT2** | Down | Glutathione metabolism | 5.3 | 4.09E-03 |
| **TermvT2** | Down | Glycerophospholipid metabolism | 5.6 | 7.03E-03 |
| **TermvT2** | Down | Glycine, serine, alanine, and threonine metabolism | 5.5 | 2.97E-03 |
| **TermvT2** | Down | Histidine metabolism | 3.5 | 2.00E-02 |
| **TermvT2** | Down | Lysine metabolism | 5.9 | 2.97E-03 |
| **TermvT2** | Down | Methionine and cysteine metabolism | 8.5 | 8.44E-04 |
| **TermvT2** | Down | N-glycan synthesis | 6.3 | 1.59E-03 |
| **TermvT2** | Down | NAD metabolism | 5.8 | 3.65E-03 |
| **TermvT2** | Down | Nucleotide interconversion | 4.7 | 1.40E-02 |
| **TermvT2** | Down | Pentose phosphate pathway | 3.7 | 3.49E-02 |
| **TermvT2** | Down | Phenylalanine metabolism | 4.9 | 4.15E-03 |
| **TermvT2** | Down | Protein degradation | 8.6 | 8.44E-04 |
| **TermvT2** | Down | Pyrimidine catabolism | 6.6 | 8.44E-04 |
| **TermvT2** | Down | Pyrimidine synthesis | 7.1 | 8.44E-04 |
| **TermvT2** | Down | Selenoamino acid metabolism | 6.3 | 2.97E-03 |
| **TermvT2** | Down | Sphingolipid metabolism | 8.0 | 8.44E-04 |
| **TermvT2** | Down | Transport, extracellular | 8.0 | 1.02E-03 |
| **TermvT2** | Down | Transport, golgi apparatus | 6.6 | 1.59E-03 |
| **TermvT2** | Down | Transport, lysosomal | 6.7 | 1.02E-03 |
| **TermvT2** | Down | Transport, mitochondrial | 4.6 | 1.40E-02 |
| **TermvT2** | Down | Tryptophan metabolism | 4.8 | 2.97E-03 |
| **TermvT2** | Down | Tyrosine metabolism | 6.3 | 1.19E-03 |
| **TermvT2** | Down | Urea cycle | 7.8 | 8.44E-04 |
| **TermvT2** | Down | Valine, leucine, and isoleucine metabolism | 4.0 | 1.64E-02 |
